# A common gene drive language eases regulatory process and eco-evolutionary extensions

**DOI:** 10.1101/2020.02.28.970103

**Authors:** Prateek Verma, R. Guy Reeves, Chaitanya S. Gokhale

## Abstract

Synthetic gene drive technologies aim to spread transgenic constructs into wild populations even when they impose organismal fitness disadvantages. The extraordinary diversity of plausible drive mechanisms and the range of selective parameters they may encounter makes it very difficult to convey their relative predicted properties, particularly where multiple approaches are combined. The sheer number of published manuscripts in this field, experimental and theoretical, the numerous techniques resulting in an explosion in the gene drive vocabulary hinder the regulators’ point of view. We address this concern by defining a simplified parameter based language of synthetic drives. Employing the classical population dynamics approach, we show that different drive construct (replacement) mechanisms can be condensed and evaluated on an equal footing even where they incorporate multiple replacement drives approaches. Using a common language, it is then possible to compare various model properties, a task desired by regulators and policymakers. The generalization allows us to extend the study of the invasion dynamics of replacement drives analytically and, in a spatial setting, the resilience of the released drive constructs. The derived framework is available as a standalone tool. Besides comparing available drive constructs, our tool is also useful for educational purpose. Users can also explore the evolutionary dynamics of future hypothetical combination drive scenarios. Thus, our results appraise the properties and robustness of drives and provide an intuitive and objective way for risk assessment, informing policies, and enhancing public engagement with proposed and future gene drive approaches.

## Introduction

Gene drive techniques increase the frequency of a synthetic genetic element in populations in a manner only partially determined by its impact on organismal fitness (and stochastic events). Swift progress in molecular biology allows us to design complicated drive systems which may be substantially more efficient in the properties of interest than their natural counterparts. The need for theoretical sandboxing of such technology with planetary consequences is imperative before field deployment. It is also critically important to provide the stakeholders of such a technology sufficient understanding to evaluate the basis of crucial projected outcomes. However, the number of publications on theoretical and experimental synthetic gene drive systems is overwhelming and ever-increasing. Generally, the description of properties of each of the sequentially proposed synthetic drive approaches uses bespoke modelling frameworks [Davis et al., 2001, Ward et al., 2011, Unckless et al., 2017]. The ability to quickly compare the relative sensitivity of the fundamental properties of different drive scenarios to parameter changes is currently limited. Regulators, policymakers, and non-experts alike desire such a tool to discuss the applicability of synthetic drive constructs. Consequently, we suggest a common language and demonstrate its applicability in a simplified framework.

The natural Segregation Distorter (SD) locus in *Drosophila melanogaster* imposes an enormous organismal fitness cost, in that it is homozygous lethal (and only viable as heterozygotes) [Sandler et al., 1959, Sandler and Golic, 1985, Crow, 1991]. Natural selection, therefore, at the organismal level, would act to eliminate the SD allele. However, because of its capacity to bias the production of SD functional sperm in +/SD heterozygotes, the allele has rapidly increased to an equilibrium frequency of 1-5% in most natural populations worldwide [Hartl, 1975, Hiraizumi and Thomas, 1984, Brand et al., 2015]. This natural system illustrates how gene drive elements can increase in frequency despite a substantial cost to (overall) organismal fitness. Developing analogous synthetic drive elements to push linked genes into wild populations in a self-perpetuating manner is an old aspiration [Craig et al., 1960]. Based on the intended use, the synthetic gene drive system can be categorized into two types: replacement drives (also modification drives) and suppression drives. Suppression drives aim to reduce or completely eradicate the target populations upon release and remain the focus of many regulatory considerations [Warmbrod et al., 2020, Moro et al., 2018]. A replacement drive works by incorporating or substituting a target gene with the desired gene into the population. Replacement drives have broad applicability ranging from the control of disease vectors by rendering them harmless [Collins and James, 1996, Isaacs et al., 2011, Gantz et al., 2015] to making resistant pests sensitive to insecticides [Collins, 2018] and control of invasive species in agriculture [Courtier-Orgogozo et al., 2017, Buchman et al., 2018]. For instance, Gantz et al. 2015 have provided evidence of a CRISPR based gene-drive system that can spread antimalarial genes into a target vector population of *Anopheles stephensi* and renders them resistant to the human malaria parasite *Plasmodium falciparum* [Gantz et al., 2015]. In agriculture, Buchman et al. 2018 have reported the construction of a synthetic Medea gene drive for a significant crop pest *Drosophila suzukii* rendering them harmless to the target crops [Buchman et al., 2018].

Like SD, it does not necessarily follow that any synthetic drive element will increase in frequency to the extent that it displaces all wildtype alleles initially present in the wild population. This fixation property is dependent on various drive parameters of the developed system. Other such properties of interest are the speed of action, reversibility, and potential to be spatially confined to only target populations. The sensitivity of such fundamental properties of drive systems to drive parameters has been a topic of interest of numerous recent theoretical studies [Huang et al., 2011, Akbari et al., 2013, Vella et al., 2017, Eckhoff et al., 2017, Noble et al., 2018, Edgington and Alphey, 2018, Dhole et al., 2018, Edgington and Alphey, 2019, Holman, 2019, Champer et al., 2020c]. We have collated this material in the provided database.

We constructed a representative literature database on synthetic gene drive system to be cognizant of the current trends in this rapidly growing field of research GitHub. The database consists of 75 publications from the year 1995 to 2021. The literature is sorted based on gene drive type (replacement or suppression), the model system under study, theoretical methodology, consideration of breakdown of drive, the possibility of gene drive reversibility and public accessibility of the literature. From the analysis of the literature database, we found that the number of studies on replacement drives is 35 [Gantz et al., 2015, Marshall and Akbari, 2016] and suppression drives are 37 [Hammond et al., 2016, Beaghton et al., 2017, Kyrou et al., 2018], with twelve publications considering both approaches. The majority of research studies (41 out of 62 total) have considered resistance evolution in synthetic gene drive systems [Unckless et al., 2017, Noble et al., 2017]. Analytical methodologies mainly employed deterministic and stochastic models with a few including spatial features [Huang et al., 2011, Eckhoff et al., 2017, Tanaka et al., 2017, Dhole et al., 2018, Girardin et al., 2019, Bull et al., 2019, Champer et al., 2019]. The model organisms in gene drive studies have been chiefly mosquitoes (total 25) [Windbichler et al., 2011, Gantz et al., 2015, Hammond et al., 2016], fruit flies (total 13) [Larracuente and Presgraves, 2012, Gantz and Bier, 2015, Buchman et al., 2018], rodents (total 3) [Lindholm et al., 2013, Grunwald et al., 2019] and 20 generic studies with no particular species in mind.

Most of the studies, we observe, use new terminologies for the bespoke drive mechanisms developed therein. It is partially because the molecular mechanisms of different gene drives can be very different and complex. While excellent molecular biology tools are used in ingenious ways to develop new synthetic drives, they still act on population genetics’s fundamental forces. The new jargon sometimes is unnecessarily confusing for policymakers and regulators in charge of deciding about such techniques’ future applicability. Since an exact comparison between the techniques is not possible, a new and useful technique might get lost in the plethora of synthetic drive projects.

The linguistic challenges in discussing the unavoidably complex gene drive topic lead to at least two types of problems. First is the inconsistent application of terminology; this can add to the confusion, and in some instance, can actively contribute to misunderstandings. For example, the use of different terms to describe the same thing, e.g. the term “quorum” in daisy quorum drive [Min et al., 2017] appears to be substantially or entirely synonymous with the much earlier and commonly used term underdominace [Davis et al., 2001]. Other examples include the equivalence between modification drive, replacement drive and population replacement or between driving-Y and non-autosomal X-shredder. Furthermore, some terms are used inaccurately as being synonymous when they are not; an example is that gene drive is frequently equated with “non-Mendelian” or “super-Mendelian” inheritance. However, this is not necessarily the case, e.g. for gene drives like Medea or underdominace where zygotes’ genotypes can be entirely Mendelian. In this light, it is notable in the recent attempt to standardise the definition of gene drive [Alphey et al., 2020], where gene drive is stated to work by “reducing the fitness of alternative genotypes without directly distorting Mendelian inheritance”.

The second source of linguistic challenges is inconsistent parametrisation and their description. This disparity almost inevitably results from the bespoke modelling approaches currently employed to describe each drive approach. To reduce, but certainly not eliminate, the above challenges, we propose terminology that permits a common parameterised synthetic drives language (using a relatively small number of standard parameters rooted in standard population genetics). This standardisation enables non-experts to precisely and quantitatively discuss a wide range of replacement drive scenarios in a mutually comprehensible manner. Furthermore, it also greatly facilitates the intuitive description of drive systems that combine multiple distinct drive approaches. This feature could be of increasing value in the future as drives get complicated. As one of the most advanced assembled drive systems, with the acronym SDGD, is a combination of two distinct drive approaches (SDGD is a suppression drive and as such is not considered in this study, which focuses exclusively on replacement drives) [Simoni et al., 2020]. Furthermore, a modelling study focused on SDGD speculates on adding a third component to this drive system to carry an anti-parasite gene [North et al., 2020]. Other modelling studies have focused on combined drives to enhance desirable gene drive properties [Gokhale et al., 2014, Faber et al., 2021, Oberhofer et al., 2019, 2020, Dhole et al., 2019, Noble et al., 2019, Min et al., 2017, Edgington et al., 2020, Willis and Burt, 2021, Champer et al., 2021, 2020a], a trend likely to accelerate in the future. Such developments considerably increase the complexity of discussing gene drives compared to single drive systems’ already existing substantial complexity.

Analyzing the select literature, we have distilled the primary components of synthetic gene drive models. From the principles of standard population genetics, we incorporate the processes that subvert the generally dominant role that organismal fitness plays in how natural selection can impact the frequencies of alleles within natural populations. Precise accounting of a generic diploid organism’s lifecycle through the various stages of development, from an adult, forming gametes to zygote and then back to an adult, is done. We discuss how the drive can act at any one or all of these stages. We then proceed to combine the knowledge into a single population dynamic model. Our model is backwards compatible, as demonstrated by the recovery of specific gene drives discussed in previous theoretical and experimental studies. Furthermore, the explicit use of standard terminology also allows us to extend the same basic model to complicated scenarios such as multilocus and multiallelic drive mechanisms (in section Backward compatibility). We deploy the developed succinct theoretical model for a single locus system in a user-friendly tool `DrMxR` - Drive Mixer (ShinyApps).

While our basic model is available as a standalone tool, we provide results also extending to an ecological and spatial dimension. A mechanism for localizing the gene drive to a target population is the imposition of a suitably high invasion threshold. We determine the extended conditions required for the invasiveness of drive. We also evaluate the impact of spatial structure on the condition of invasion (from rare) and fixation of the drive for a single population.

When considering multiple proposed drive systems for release, regulators find it essential that a comparison between the systems is possible. The current state of the art does not allow this easily. We thus show that a single theoretical approach, when minimally extended, provides specific cases of different drive systems. This exercise provides us with a common vocabulary across different drive systems. Furthermore, we provide `DrMxR` to test specific cases of proposed drive mechanisms that will be useful in risk assessment and regulation of gene drives. With applicability in mind, `DrMxR` is specially targeted towards policymakers, the general public, and even experts for quick hypothesis testing. To this end, we begin by detailing the process of theory development in the following section.

## Results

For developing this model, we have assumed an obligate sexually reproducing organism, a likely necessity for successful gene drive where organismal fitness is negatively impacted. We split the life cycle of an organism into three tractable stages; the minimal abstraction required to recover the established results in the field of engineered gene drive systems. Further complications can indeed be added depending on the details of the case study in focus.

Figure 1 shows the lifecycle of an individual in our model. We focus on two allelic types - wildtype (W) and the driven gene (D). Thus we have adults of three genotypes, wildtype homozygotes WW, heterozygotes WD and drive homozygotes DD. Adults are chosen from the population pool for reproduction. Adults produce gametes that combine to form zygotes. The zygotes grow up to become adults, and the cycle continues. We allow for overlapping generations, a realistic assumption for numerous target species such as mosquitoes, drosophila or rodents [Windbichler et al., 2011, Backus and Gross, 2016, Buchman et al., 2018]. We assume that the alleles during gamete formation are segregated independently according to Mendel’s inheritance laws. Hence the total number of alleles in the absence of any evolutionary processes remain conserved over successive generations. Therefore, frequencies of genotypes reach Hardy-Weinberg equilibrium in the limit of an infinite population, random mating, and no selection.

**Figure 1:**
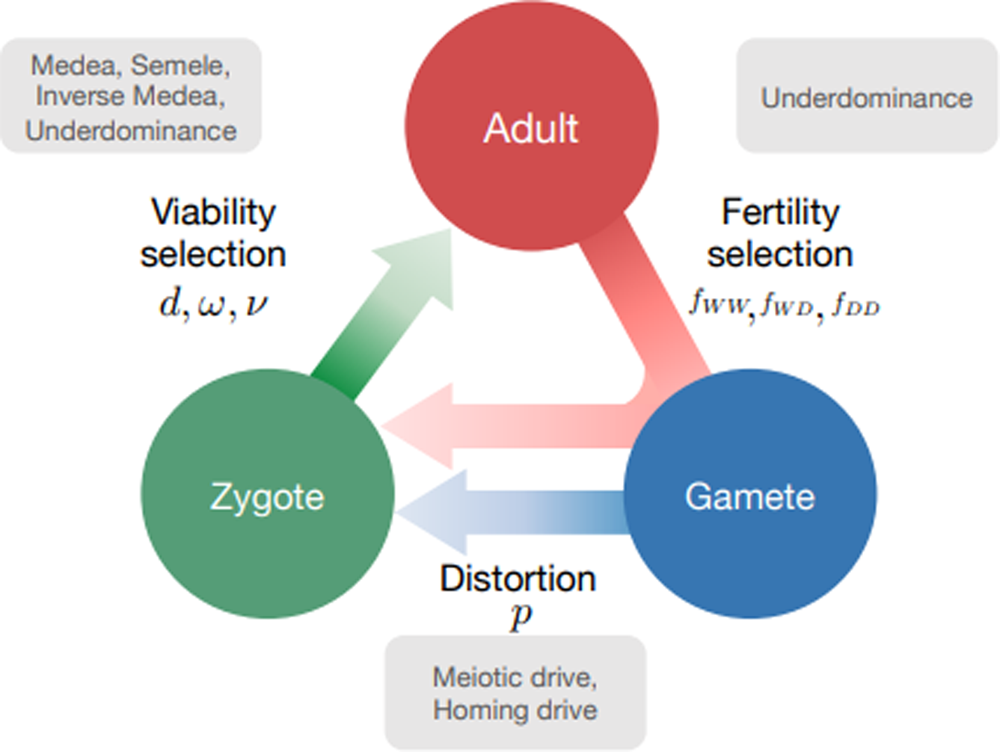
Lifecycle of an individual organism for a generic gene drive model. Assuming that individuals reproduce sexually and that the lifecycle has three stages, Adult, Gamete and Zygote. Adults produce gametes which combine to form zygotes. Zygotes grow up to become adults. Three factors can act during the life stages of an organism: distortion, viability selection and fertility selection (represented as arrows). Each can influence the probability of inheritance of a gene in the population and can be potentially manipulated to engineer gene drive constructs. Parameters, described in the text, are associated with each of the three arrows. Examples of named drive systems that can be generated are provided associated with the respective arrow.

The essential feature of a gene drive is biasing the chance of inheritance of the desired gene in the population [Champer et al., 2016]. The expected outcome, however, is that the population composition is modifiable in a controlled fashion. Interventions along the lifecycles can accomplish the change via distortion, viability and fertility selection. These processes act at different stages of an individual’s life cycle. Distortion acts at the gamete level and biases the transmission of the drive allele in the heterozygote. Gametes combine to form zygotes, but some are non-viable and die. Fertility selection acts at the adult stage when individuals are chosen to reproduce with probability proportional to their fitness. Distortion, viability selection and fertility selection, thus, together or even independently, can drive the population away from the Hardy-Weinberg equilibrium. Synthetic gene drive techniques allow us to engineer such selection pressures.

### Viability selection

Viability selection acts during the zygote phase of an individual’s lifecycle. The viability fitnesses represent the inherent variation in the fitnesses of the three genotype, WW, WD and DD. The fitness can also capture the payload costs of the drive allele. Viability fitness is defined here as the probability of survival of the zygotes up-to-the adult stage. *ω* and *ν* denotes the genotypic viabilities of WD and DD, respectively. The above parameters have been normalized to the viability of WW fixed at 1.

Well described synthetic drive systems that work principally by manipulating viability selection parameters include those using zygotic toxin-antidotes. In these systems, a proportion of zygotes of specific genotypes may become non-viable. Medea (Maternal effect dominant embryonic arrest) is an example of a naturally occurring toxinantidote gene drive found in flour beetles [Beeman et al., 1992, Wade and Beeman, 1994]. In Medea drive, wildtype homozygous offspring of heterozygous mothers are non-viable. Population dynamics of Medea drives have been studied in Ward et al. [2011], Gokhale et al. [2014]. A synthetically engineered Medea drive first demonstrated in *Drosophila* Chen et al. [1997] has been extensively studied Akbari et al. [2014], Buchman et al. [2018]. Similarly, a synthetic viability selection based underdominant population transformation system was developed for *Drosophila melanogaster* in Reeves et al. [2014]. Figure 2A shows the population dynamics of Medea drive and deviation from Hardy-Weinberg equilibrium parabola.

**Figure 2:**
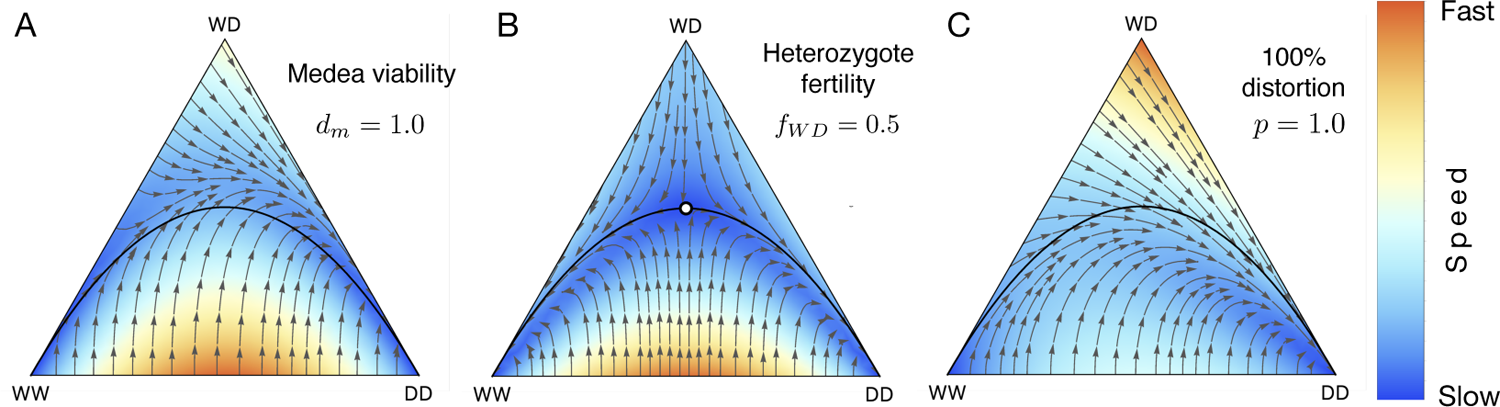
Effects of fertility selection, distortion and viability selection on population dynamics of the three genotypes. Population consists of single genotype at the vertices of a triangle in de Finetti diagram. A point in the interior corresponds to the population composition where all three of the genotypes potentially exist. Their relative abundance is proportional to the distance from the vertices. The black parabola curve represents Hardy-Weinberg equilibrium. The white open point represents the population composition of the fixed point. Colours exhibit speed of the dynamics inside de Finetti plots. The speed of the dynamics has been normalized for each plot and their absolute values are not directly comparable between diagrams. **(A)** Viability selection for Medea gene drive with drive efficiency *dm* = 1. **(B)** Fertility selection for the underdominance case where fertilities of the genotypes are *f_WW_* = 1, *f_WD_* = 0.5, *f_DD_* = 1. An unstable point appears in the interior of de Finetti diagram and is denoted by a white circle at (*x_WW_, x_WD_, x_DD_*) = (0.25, 0.50, 0.25). **(C)** Distortion when drive heterozygous individuals contribute drive allele with 100% efficiency i.e. *p* = 1.

The result can be recapitulated by readers using `DrMxR` (ShinyApps) where Medea and other related synthetic drive systems can be seamlessly modelled including inverseMedea [Marshall and Hay, 2011], or Semele [Marshall et al., 2011]. The drive efficiencies of Medea, Inverse Medea and Semele drive is represented by parameters *d_m_*, *d_im_* and *d_s_* respectively. We recover a subset of key results of the population dynamics from earlier publications in the backward compatibility section. The framework used by `DrMxR` is general and applicable to other single construct gene drive systems either entirely or partially based on viability selection.

### Fertility selection

Specific genotypes may experience fitness advantages because of preference for traits during mating or because some genotypic pairings are more fertile than others. Both of these fitness components are modelled using the fertility selection parameters. The fact that both mating success and fecundity are considered jointly dictates that the fertility selection arrow on Figure 1 traverses three life stages, rather than the two indicated for the other types of selection. The fertility fitness component arising from mating success is included in the parameter *f_WW_*, *f_WD_* and *f_DD_* for the three genotypes. Fertility selection is an evolutionary phenomenon that drives the population away from the Hardy-Weinberg equilibrium. Our model did not differentiate between sexes, but it is possible to include this complexity [Hofbauer and Sigmund, 1998].

Previous work [Feldman and Liberman, 1985, Nagylaki, 1987] captures the rich dynamics that ensue when fertility selection is considered. The population dynamics of a two allele system for different fertilities and sex-dependent viabilities have been extensively studied in Hofbauer and Sigmund [1998]. The authors have also accounted for non-random mating between the mating pairs by introducing additional parameters [Hofbauer and Sigmund, 1998]. We have accounted for variable fertility rates by introducing suitable parameters in the context of the gene drive system (as shown in Figure 2B).

### Distortion

Gametic distortion alters the transmission of drive alleles in heterozygotes, so they substantially exceed the Mendelian expectation of 50% and is controlled by the single parameter *p* in our model. Biologically such distortion happens in natural meiotic drives where meiosis is subverted due to intra-genomic conflict [Sandler and Novitski, 1957, Palopoli and Wu, 1996, Lindholm et al., 2016]. Examples of naturally occurring gene drive elements based on distortion are segregation distorter and t-haplotype in heterozygous fruit flies and mice, respectively [Lyon, 2003, Larracuente and Presgraves, 2012]. These drive elements bias their transmission during spermatogenesis by killing sperm carrying non-driving alleles (W). Though the killing of non-carrier sperm also has the potential to reduce fertility [Price and Wedell, 2008, Lindholm et al., 2016], ‘distortion’ can be conceived as an independent evolutionary force responsible for biased transmission of drive allele. The synthetic homing drive also distorts the transmission of alleles in heterozygotes. To keep the model tractable, both analytically and in terms of user comprehension, `DrMxR` does not currently consider sex-ratio gene drives (Y-driving, X-Shredder) [Burt, 2003, Burt and Deredec, 2018]. Figure 2C shows the effect of distortion on the population dynamics of the three genotypes: WW, WD, DD. Previously published evolutionary dynamics of a homing drive using CRISPR are recovered in the Methods section.

All the above methods of biasing the inheritance pattern of a gene are recovered employing our generic model. In Methods, we first derive the mathematical formulations of the processes independently and then combine them in a single dynamical model system. To demonstrate the generality of our approach, we recover the results of Noble et al. [2017], Marshall and Hay [2011], Marshall et al. [2011] and Gokhale et al. [2014] as special cases of our model formulation. Ecologically it is vital to characterize the spread of a genetic construct. We do this in panmictic as well as spatially constrained populations. We provide an analytical form for calculating the refraction zone (the safe amount of drive heterozygotes and homozygotes from which the wildtypes can recover). For spatially constrained systems, we show the exact form in which the probability of invasion and fixation of a drive element depends on the network’s connectivity.

### Combined dynamics

The three factors viz. distortion, viability and fertility selection can act during the three stages of an organism’s lifecycle. Figure 2 illustrates the specific impacts of these forces on the population dynamics by varying parameters using our application `DrMxR`. The equilibrium dynamic changes in different ways relative to the Hardy-Weinberg equilibrium line in Figure 2. Besides individual impact, our application allows intuitive exploration of scenarios when more than one of these three evolutionary forces acts in combination. Realistically, such scenarios arise when a drive element impacts simultaneously both distortion and fertility selection [Price and Wedell, 2008, Lindholm et al., 2016]. In the Drosophila segregation distorter, selective killing of sperm carrying a wildtype allele in heterozygous males biases the transmission of drive allele and potentially reduces the males’ fertility. Homing endonuclease gene drives based on CRISPR/Cas9 have been mathematically modelled to bias transmission and also to reduce the fertility of the genotype carrying payload gene [Noble et al., 2017].

Our approach recovers the result of Noble et al. [2017] showing the combined effects of distortion and fertility selection on population dynamics. Additionally, our application allows us to study various drive combinations as well. In the Methods, we recover the result of Gokhale et al. [2014] and show the combined effect of fertility selection (underdominance) and viability selection (Medea gene drive). Similar explorations of the population dynamics of other drive combinations across their entire parameter range are possible in `DrMxR`, for example, Medea (viability selection) together with homing endonuclease (distortion) can be studied.

### Ecological factors

In the context of field deployment, understanding only the population genetics of the system is not enough. The properties of gene drive constructs are diverse, depending on their molecular construction and the differential selection pressure they impose in the varied ecological situations. Conversely, the ecology of the target species it-self can disrupt the intended dynamics of the driven gene. Taking the demographic parameters into account is imperative when assessing the impact of gene drive deployment. Below we derive the invasion threshold of a drive system and evaluate the impact of spatial structure on the invasion (from rare) and fixation of the drive for a single population.

### Invasion threshold

The unintended spread of certain types of drive to non-target populations has been a significant concern ever since the conception of synthetic gene drives. This interest is particularly the case for replacement drives (not intended to alter the size of populations) since the negative selection costs (fertility and viability) imposed by replacement-drive constructs are generally much smaller than for suppression drives [Dhole et al., 2018, Edgington and Alphey, 2018, Marshall and Hay, 2012, Ward et al., 2011]. In this context, the option of localizing the replacement gene drive to target populations has been the focus of scientists for both developing and regulating gene drive [Backus and Delborne, 2019]. A mechanism for localizing the driven construct is the imposition of a suitably high invasion threshold. The invasion threshold is the minimum frequency of drive carrying organisms required to be released to replace the wild target population. If the invasion threshold is high, the drive is likely more spatially restricted because the invasion of the non-target populations will require a large number of introduced individuals. As high threshold drives theoretically limit their spatial spread, they also may mitigate the spread of drives into partially interfertile species (or subspecies). Accidental release of a few drive organisms may completely transform wild populations for gene drives with low or no threshold [Marshall, 2009, Noble et al., 2018]. A recent review of different types of gene drives based on a quantitative analysis of their invasiveness can be found in Frieß et al. [2019].

A relevant quantity of interest is the number of drive individuals required to invade a wild population successfully. Here we consider both the drive heterozygotes and homozygotes together as drive individuals. In our model, the invasion threshold can be quantified based on the direction of the flow lines in the de Finetti diagram. We define the refractory zone as the area of the flow lines towards the population consisting of all wildtypes in the de Finetti diagrams. Thus, we quantify the amount of release that a population may sustain and still revert to the wildtype by measuring the wild-type vertex’s basin of attraction.

We calculated the refractory zone by analytically computing the equation of the invariant manifold separating the flow lines through approximations. Details of the calculation are in the Methods section. The refractory zone quantifies the minimal number of drive heterozygotes and homozygotes (released or migrants), capable of transforming the wildtype population.

In the model, variation in the drive efficiency and fitness of different genotypes affects the refractory zone of a gene drive system. Using the insight provided from Fig. 1, we consider the case of distortion based gene drive along with fertility selection. Figure 3 shows the heat-map of the refractory zone with variation in distortion probability *p* and fertility fitness of heterozygotes *f_WD_*. When both the drive efficiency and fitness of heterozygous are high, the distortion drive’s refractory zone is zero. Hence an accidental release of only a small frequency of drive organism would lead to complete replacement of the wild population. The gene drive system is, therefore, absolutely non-localized. Low distortion drive efficiency and fitness of heterozygotes make the drive system localized, so a significant release of drive organism is required to successfully transform the wild population [Tanaka et al., 2017]. For intermediate values of *p* and *f_WD_*, the drive system is localized and does not require a massive release [Tanaka et al., 2017, Deredec et al., 2008].

**Figure 3:**
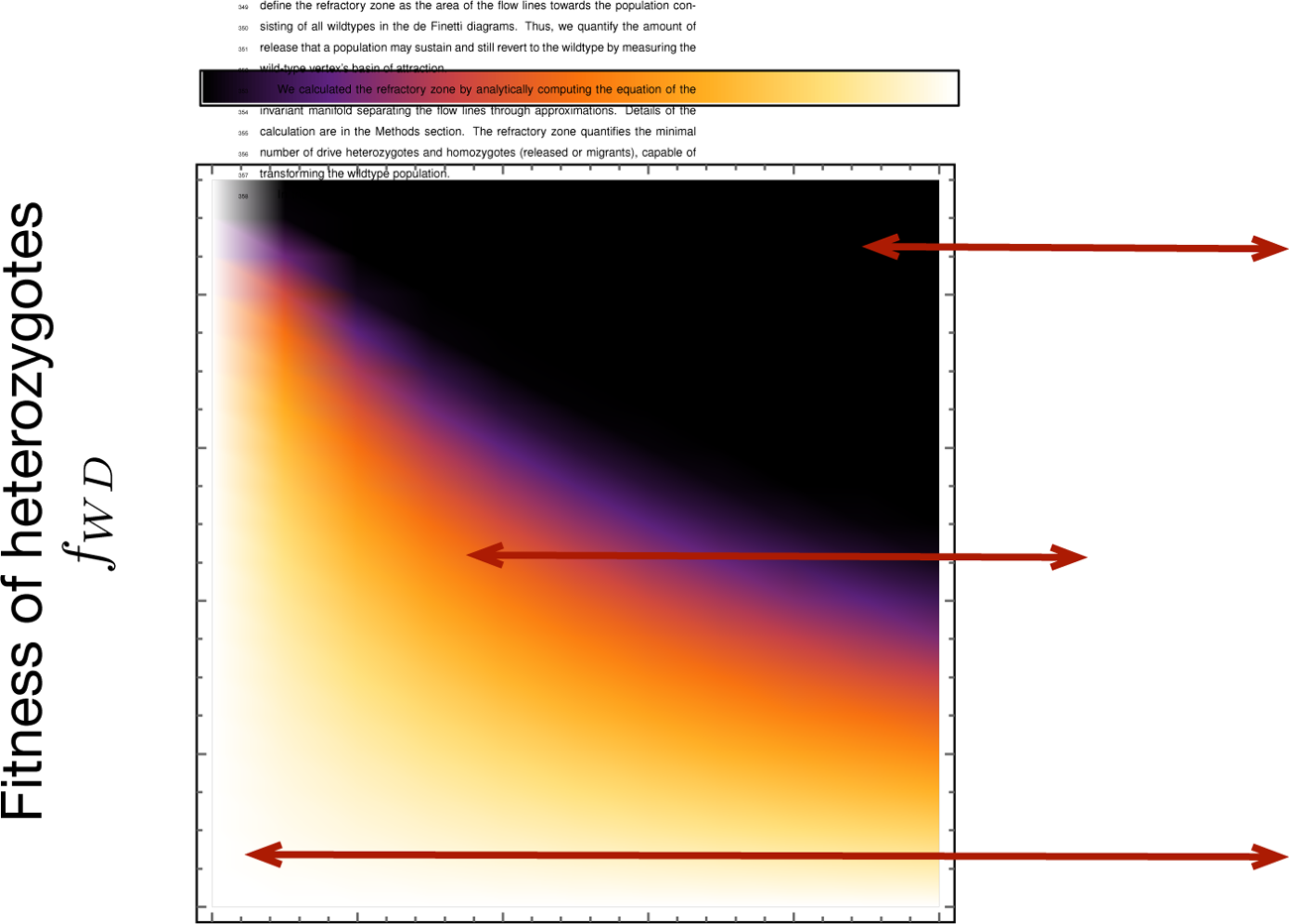
Heat-map showing the refractory zone with variation in distortion probability *p* and fertility fitness of heterozygotes *f_WD_*. Illustration of refractory zone for specific values of *p* and *f_WD_* of the heat-map. Trajectories of a de Finetti diagram when 2*pf_WD_ > f_WW_*, drive individuals invade the wild population. Refractory zone is zero and is shown by black colour in the heatmap. *p* = 0.5 corresponds to ‘no distortion’ case. The values of other parameter is fixed to *f_WW_* = 1, *f_DD_* = 1.

### Spatial organization within a population

Recent works have highlighted the need for realistic spatial modelling for more accurately predicting the outcome of gene drive release more so for suppression [Huang et al., 2011, Champer et al., 2019, North and Godfray, 2017, North et al., 2019, 2020, Tanaka et al., 2017, Eckhoff et al., 2017] than the replacement drives [Girardin et al., 2019]. Most of the analytical models, including `DrMxR`, assume random mating between individuals of different genotypes. Nevertheless, assuming random mating may give an incorrect prediction about the invasion condition of the gene drive. In reality, individuals in the population are spatially constrained and more likely to interact with individuals living in proximity. This factor will interfere with the evolutionary dynamics of the spread of gene drives. Consequently, we have developed a framework to explore the consequences of relaxing the assumption of a well-mixed population. Here we derive the condition for a distortion based gene drive to invade a single wild population if the assumption of random mating is violated and the population is spatially structured. The details of this derivation are given in the Methods section.

The analysis (See method section) uses the framework of evolutionary game theory and tracks the frequencies of alleles instead of genotypes. Previous work has shown that interpreting the association of alleles in a diploid genome as a two-player game leads to some intriguing new insights into genetic evolution [Hofbauer et al., 1982, van Veelen, 2007, Traulsen and Reed, 2012]. Also, different ways of updating a population can lead to different allele dynamics in a panmictic population. [Traulsen et al., 2006, 2005]. Population update rule defines the elementary process that changes the frequency of each type in the population; for example, in the birthdeath update rule, an individual is selected first for birth proportional to its payoff from the evolutionary game. It replaces another randomly chosen individual from the population selected for death. In our case, population update occurs in allele space, so an individual unit is an allele that can be wildtype (W) of drive type (D).

Ohtsuki and Nowak [2006] found that if the interaction between the players (alleles in our case) take place on a regular graph of degree *k* (see figure 4), the payoff entries of the game are transformed according to equation (13). So, as *k* tends to infinity, the additional transformational entries of payoff matrix will become increasingly small, and invasion of gene drive will essentially depend upon whether the drive allele is more fit than wildtype when the drive is rare, and fixation will depend on whether the same is true when wildtype is rare. Since the interaction unit in this formulation is at the allele level, the biological interpretation of *k* is not straightforward. Intuitively, parameter *k* measure the level of mixing between individuals within a population (where *k* tending to infinity corresponds to complete mixing, a simplifying assumption common to many models including `DrMxR`). Since a different mathematical formulation has been employed, results obtained in this section cannot be added to the `DrMxR` where dynamics can be visualised through de Finetti plots.

**Figure 4:**
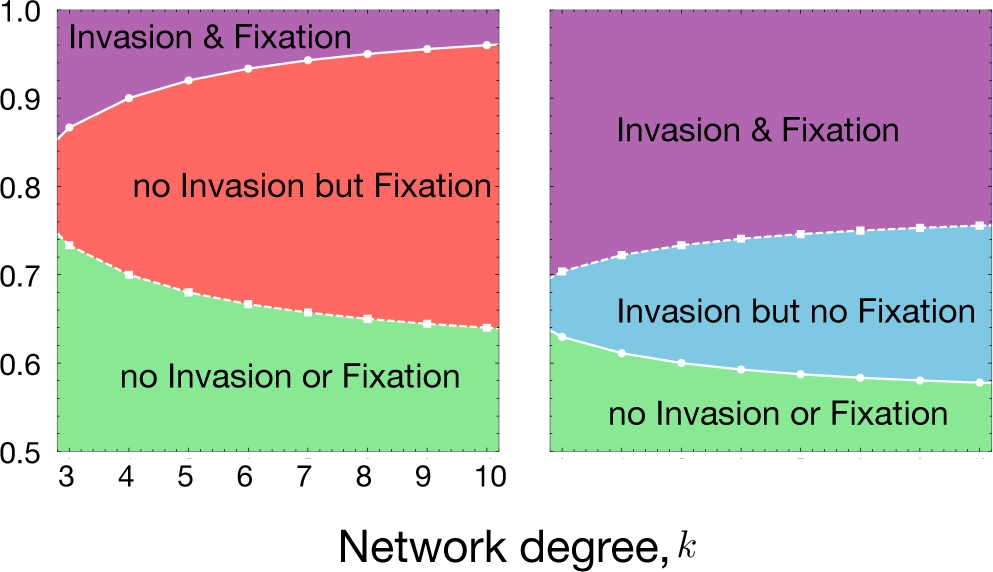
Spatial structure affects the condition for the invasion from rare and fixation of the driven gene. **(A)** Variation in invasion (full line with circles) and fixation (dashed line with squares) conditions with respect to network degree (*k*) and distortion parameter (*p*) for *f_WD_* = 0.5 and **(B)** *f_WD_* = 0.9. The values of other parameters are fixed to *f_WW_* = 1, *f_DD_* = 0.4. Population dynamics changes when the population becomes more structured on the Bethe lattice parameterized by *k*. Lower *k* means more structured population and higher *k* represents less structure (closer to well-mixed case). The change in population dynamics properties can be seen by the change in invasion/fixation condition and combinations of them, such as no invasion from rare but fixation, if sufficient drive individuals are released/migrate.

Figure 4 shows that the invasion and fixation outcomes within a single population vary depending on the degree of spatial mixing and distortion efficiency. Increasing network degree can move a population where the drive cannot invade or fix to a situation where the drive can fix but cannot invade from rare for lower to moderate values of *p* (*p* = 0.65 to 0.80). The fixation but no-invasion case corresponds to the introduction of the invasion threshold that can help local confinement of the gene drive. Interestingly, one can move to this regime by regulating the degree of the network. For higher values of *p >* 0.80 when the drive can both invade and fix in the population, increasing the network degree can introduce an invasion threshold. A similar trend ensues in Figure 4B, but increasing network degree may allow the drive to invade the wild population but does not allow it to get fixed in the population. This scenario corresponds to the over-dominance case, and mathematically, the dynamics correspond to a stable fixed point in the interior of the simplex. The condition for the fixation and the invasion tends towards a well-mixed population regime for higher *k*.

## Discussion

We have developed a minimalist modelling framework and identified three forces/factors responsible for propagating gene drive in the presence of an organismal fitness cost. These forces act during different stages of the target organism’s lifecycle and relate the gene driving mechanism to the organism’s biology. Such a type of approach is arguably missing in earlier works on gene drive. For example, Noble et al. [2017] studied the population dynamics of CRISPR gene drive without explicitly stating that the fitness they incorporated belongs to fertility selection parameters. In other models fitness costs have been introduced through viability fitness parameter [Marshall and Hay, 2011, Marshall et al., 2011, Gokhale et al., 2014]. With our approach, we can highlight that the evolutionary outcome for the two cases (drive acting through viability or fertility but leading to similar costs) differs substantially. Our work stresses the importance of both the target organism’s biology and knowing the exact phases of the lifecycle where the synthetic construct will act. The current modelling approach also provides a classification of a simple gene drive system based on the biology of how the drive is designed (out of the three primary life stages) and avoids unnecessarily new and confusing terminology.

As with different translational evolutionary biology applications, the eventual aim of several synthetic gene drive constructs is field deployment. Thus, any drive technology needs to be compared with other available techniques, not by experts of the particular system but regulators who need a broader perspective. Our work employs standard population genetics methods while keeping our model as generic and minimal as possible. The resulting model allows us to provide a birds-eye view of the dynamics over the space of different drive mechanisms. Educators and regulators would benefit from using our `DrMxR` for studying the population dynamics of the gene drive. Unlike SLiM, a scriptable evolutionary simulation framework not limited to drive systems [Haller and Messer, 2019], `DrMxR` is specific to drive systems and only valid for a generic species with gamete, zygote and adult life stages. On the other hand, MGDrive, an R-package focusing on testing gene drives in species with Egg-larva-pupa-adult life stages or chiefly mosquitoes [Sá nchez C et al., 2019]. In species with density-dependent larvae competition, the timing of the expression of driving endonuclease becomes very significant, i.e. before or after the density-dependent larvae competition [Godfray et al., 2017]. `DrMxR` is currently not capable of modelling such scenario. `DrMxR` is also no substitute for species and geography specific gene drive models [North et al., 2013, Eckhoff et al., 2017, North et al., 2019, 2020]. The utility of our framework lies in easing the understand of the gene drive mechanism and how it can arise or be a by-product of distortion, viability selection and fertility selection. Though not unique, our model also distinguishes the origin of the fitness cost of the drive allele. The fitness cost can affect the fertility of the organism where the transgenic grows up to reach the adult stage, or it could also affect its viability, in which case the organism dies at the zygote stage. This distinction is crucial as it leads to different population dynamics for the same amount of fitness cost.

In our model, users can choose the driving factor and its corresponding effect on the target organism’s biology by tweaking the various parameters explored in this manuscript. Deviations from the null Hardy-Weinberg equilibrium may be studied via the effect of the three driving factors, individually or combined. It is possible to investigate conditions for invasion and fixation of the drive and its tolerance to fitness cost that is highly relevant for drive deployment (relevant code provided on ShinyApps. As case studies of our approach, we have recovered the results of various drives such as CRISPR homing endonuclease drive, Medea, single-locus engineered underdominance, Inverse Medea, and Semele in the Methods section [Marshall et al., 2011, Marshall and Hay, 2011, Gokhale et al., 2014, Noble et al., 2017].

Empirical studies have shown that the selfish genetic elements based on transmission distortion can reduce both fertility (offspring production) [Dyer and Hall, 2019, Larner et al., 2019] and viability (egg to adult ratio) [Finnegan et al., 2019] of the target species. To estimate the evolutionary outcome, we have allowed to jointly vary the factors influencing the propagation of such gene drives. Flexibility to see the combined effect for various evolutionary factors influencing the spread of gene drive on the population dynamics is an essential feature of the `DrMxR`. We believe that an-alytical results for evaluating the refractory zone would help regulators estimate the drive’s invasiveness. Methodologically, the refractory zone calculation is a development deriving from a dialogue between evolutionary games and population genetics [Altrock et al., 2010, Traulsen and Reed, 2012].

Our results show how gene drive invasion and fixation conditions differ relative to the mixed population model. We found that for lower values of network degree, the region of phase space in figure 4 for invasion & fixation and no invasion or fixation increases. Hence, introducing spatial features during interaction makes the drive either highly invasive or redundant. These results might be informative for the decisionmaker in developing an intuitive understanding of how gene drive dynamics differ for structured population instead of the common assumption of well-mixed. Also, our spatial model does not help to directly compare the potential of different drives to invade a new population through migration [Gokhale et al., 2014, Noble et al., 2019, 2018, Dhole et al., 2019, 2018].

In this study, we develop a common vocabulary to model various synthetic (and natural) gene drive systems, but the mathematical model we used cannot be regarded as general. Our current model cannot address the reduction in population size and its effects on the spread of a gene drive. Therefore, DrMxR is currently only appropriate for studying gene drives that can only bring about population replacement without affecting population density. Suppression drives – intended to eradicate or reduce the target population or ‘reversal drives’ – intended to reverse the genetic alteration introduced by the first gene drive [Esvelt et al., 2014, DiCarlo et al., 2015, Vella et al., 2017, Edgington and Alphey, 2019] are not included in the app. Some newly proposed gene drive systems that are mainly intended for suppression but can also be used for replacement, such as CleavR, TARE, TADE, double-drives and Y-linked editors, cannot be currently modelled in this study [Oberhofer et al., 2020, 2019, Champer et al., 2020a,b, Willis and Burt, 2021, Prowse et al., 2019]. Classification of such complex drive system based on our mathematical model would also be problematic since these might have very complex selection mechanisms or have simple mechanisms but whose dynamics critically depend on the genetic makeup of different populations.

Self-exhausting drives that first rapidly spread in the population and then self-exhaust after limited generations are also not included in the current version of `DrMxR` app. However, a simple form of the self-exhausting drive called daisy-chain drive has been shown in the backward compatibility section as an example of how the current mathematical model could be extended [Noble et al., 2019]. Numerous drive studies have now extended to multi-locus systems, further expanding the vocabulary of the dynamics. Currently, our application (`DrMxR`) focuses on a single locus and highlights the complexities that single-locus drives can generate. Since we root our vocabulary in processes underlying multi-locus and multi-allelic drives, our concept can be extended for multi-locus and multi-allelic drive systems such as one locus two toxin (1L2T), two locus two toxin (2L2T) and reciprocal chromosomal translocation (RCT), Killer & rescue drive and tethered homing gene drive [Davis et al., 2001, Dhole et al., 2018, Champer et al., 2020c, Noble et al., 2019, Dhole et al., 2018, Champer et al., 2020c, Curtis, 1968, Champer et al., 2020c, Gould et al., 2008, Dhole et al., 2019]. We have heuristically demonstrated the extension of mathematical modelling of such systems together with resistance evolution for CRISPR homing drive in the Backward compatibility section. However, these gene drive systems are not implemented in the `DrMxR` app. These extensions will also allow for the inclusion of multiple drive systems in an ecological context in the future [Dhole et al., 2020].

An important aspect of risk assessment for regulators is the ability of a gene drive to invade non-target populations through migration [Gokhale et al., 2014, Noble et al., 2019, 2018, Dhole et al., 2019, 2018, Altrock et al., 2011]. We have extended our analysis to spatial systems using game theoretical methods as per Ohtsuki and Nowak [2006]. Studying density-dependent migrations between patches [Altrock et al., 2011, Gokhale et al., 2014] could be included to understand the spread of different drive systems. Currently, DrMxR does not model such a scenario, and it can be the probable direction of future work. Inclusion of ecological parameters such as seasonality and environmental disturbances would also be necessary when utilizing the theory to model a specific target species [Eckhoff et al., 2017]. Inclusion of ecological factors such as density dependence, spatial organization, non-random mating and target specific mating systems is in progress. It will be a necessary litmus test in assessing any drive deployment strategies [Dhole et al., 2020]. For specific species, considering detailed life history and influences in the organism’s lifecycle would be a valid extension. For example, a mosquito lifecycle consists of egg, larva, pupae and adult stages. It becomes essential to distinguish when driving endonuclease is expressed, before or after the density-dependent larvae competition [Godfray et al., 2017]. Hence, adding an appropriate life cycle depending on the model organism is necessary for a more reliable prediction of gene drive spread. However, we emphasize the disparity between the theoretical developments in simple synthetic drive scenarios and the urge towards a unified understanding at the elemental level. Using a common language will allow for a comparison between different drive techniques and adaptable to complex drive systems.

## Conclusion

The vast, diverse and growing literature in the field of gene drive is often challenging to follow for non-experts because of the varying terminology. This linguistic heterogeneity obscures actual novel results and prevents a clear view of the field. The diverse vocabulary also does not facilitate easy comparisons between different drive techniques. We develop a common vocabulary describing gene drive systems based on pre-existing standard population-genetic terminology (distortion, fertility selection and viability selection). Based on this common vocabulary, we present DrMxR, a tool to grasp different gene drives while considering ecological and evolutionary aspects. We demonstrate that our model can be used to recover work already presented in several studies. Besides comparing available drive constructs, our tool is also helpful to explore the evolutionary dynamics of future hypothetical combination drive scenarios. The results obtained for drives in spatially structured organisms could be informative in developing an intuitive understanding of how gene drive dynamics differ for structured population instead of the common assumption of panmictic population. We believe that our work will be useful for regulators, educators, the general public, and even experts in developing insights about the population dynamics of the proposed and future gene drive system.

## Methods

We consider diploid individuals of single locus with two alleles: wildtype (W) and drive allele (D). The possible genotypes are WW, WD and DD. We start with the simplest case assuming an infinitely large population, random mating, random segregation of alleles during meiosis and no distinction between male and female genotype in terms of not tracking their distinct genotype frequency. We will relax some of these assumptions as we proceed. Considering all mating pairs in table 1(a), the rate of production of each genotype can be written as:

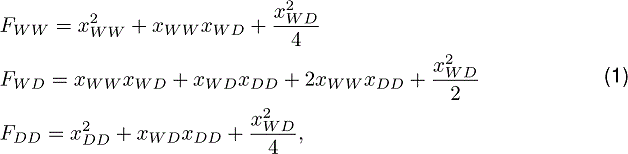

**Table 1:** Offspring proportions when alleles are segregated randomly during meiosis.

| Parents |  | Offspring |  |  |
| --- | --- | --- | --- | --- |
| ♂ | ♀ | WW | WD | DD |
| WW | WW | 1.0 |  |  |
| WW | WD | 0.5 | 0.5 |  |
| WW | DD |  | 1.0 |  |
| WD | WW | 0.5 | 0.5 |  |
| WD | WD | 0.25 | 0.5 | 0.25 |
| WD | DD |  | 0.5 | 0.5 |
| DD | WW |  | 1.0 |  |
| DD | WD |  | 0.5 | 0.5 |
| DD | DD |  |  | 1.0 |

where *x_α_* and *F_α_* are the frequency and rate of genotype production respectively and *α ∈* (WW, WD, DD). The population dynamics of the genotypes in continuous time is governed by the following set of differential equation:

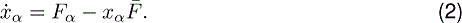

Here, *F*^-^ is the average fitness of the three genotype:

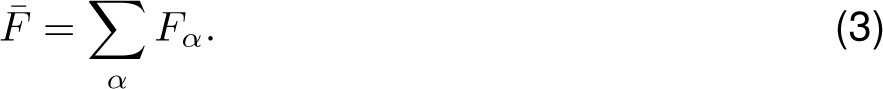

The total population remains constant hence the frequencies of all genotypes sum to unity.

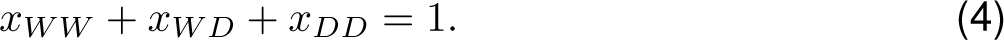

Constraints on frequencies allows us to represent the dynamics of (2) in a de Finetti diagram. We will now derive the population dynamics equations when the three factors, namely viability selection, fertility selection and distortion, are added to the system one by one.

### Viability selection

Viability selection is observed in many toxin-antidote gene drive constructs. These drives adhere to Mendel’s inheritance laws and do not distort the transmission of alleles at the gamete level. In such systems, particular offsprings become non-viable during zygote stage of the life cycle. Examples include Medea, Inverse Medea, Semele and engineered underdominance drive etc [Beeman et al., 1992, Marshall and Hay, 2011, Marshall et al., 2011]. Depending on the type of gene drive construct one can write the rate of genotypes formation as shown in table 2 and Appendix table A.1. Independent of the toxin-antidote construct, variation at the genotype level may also give rise to variation in the viabilities, that is, the probability of survival of a zygote up to the adult stage. Here *ω* and *ν* are the genotypic viabilities of the drive heterozygotes (WD) and homozygotes (DD) respectively. The rate of zygote production in the next generation for Medea, Inverse Medea and Semele gene drive can be written as:

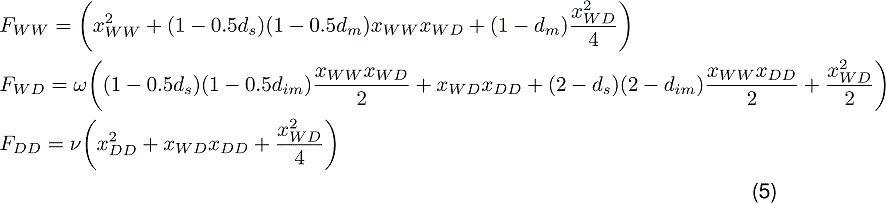

**Table 2:** Effect of fertility selection, distortion and viability selection on mating rates and offspring proportions. (a) Fertility selection changes the mating rate of genotypes at adult stage. (b) Distortion biases the transmission of drive allele from heterozygous individual by probability *p >* 0.5. Each entry in offspring’s column gives the proportion of genotype produced from the mating pair in the corresponding row. (c) Viability selection effects offspring proportions as some may become non-viable. To illustrate an example, we consider Medea gene drive where wild-type homozygous offspring of heterozygous mother are non-viable.

| Fertility Selection |  |  | Distortion |  |  | Viability Selection (e.g. Medea) |  |  |
| --- | --- | --- | --- | --- | --- | --- | --- | --- |
| ♂ | ♀ | Fertilities | WW | WD | DD | WW | WD | DD |
| WW | WW | $f_{WW} \cdot f_{WW}$ | 1.0 | | | 1.0 | | |
| WW | WD | $f_{WW} \cdot f_{WD}$ | $(1 - p)$ | $p$ | | $0.5(1 - d_m)$ | $0.5\omega$ | |
| WW | DD | $f_{WW} \cdot f_{DD}$ | | 1.0 | | | $1.0\omega$ | |
| WD | WW | $f_{WD} \cdot f_{WW}$ | $(1 - p)$ | $p$ | | 0.5 | $0.5\omega$ | |
| WD | WD | $f_{WD} \cdot f_{WD}$ | $(1 - p)^2$ | $2p(1 - p)$ | $p^2$ | $0.25(1 - d_m)$ | $0.5\omega$ | $0.25\nu$ |
| WD | DD | $f_{WD} \cdot f_{DD}$ | | $(1 - p)$ | $p$ | | $0.5\omega$ | $0.5\nu$ |
| DD | WW | $f_{DD} \cdot f_{WW}$ | | 1.0 | | | $1.0\omega$ | |
| DD | WD | $f_{DD} \cdot f_{WD}$ | | $(1 - p)$ | $p$ | | $0.5\omega$ | $0.5\nu$ |
| DD | DD | $f_{DD} \cdot f_{DD}$ | | | 1.0 | | | $1.0\nu$ |

Here *d_m_*, *d_im_* and *d_s_* measures the drive efficiency of Medea, Inverse Medea and Semele drives respectively. An example of how viability selection can be implemented is shown in the case of Medea in Table 2.

### Fertility selection

The relative number of offsprings produced from reproduction may differ because of the variation in the fertilities of the adult mating pairs. The fitness component due to differential fertilities can be incorporated in the parameters *f_α_* where *α ∈* (WW, WD, DD). The rate of the offspring production for the three genotypes because of fertility selection changes to

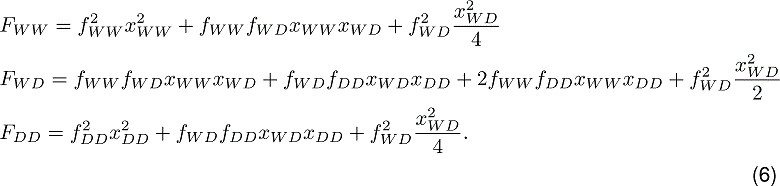

The population dynamics is again given by equation (2). We assume in equation (6) that all the offsprings have equal viabilities *ω* = *ν* = 1 and no toxin-antidote drive is present hence *d_m_* = *d_im_* = *d_s_* = 0. A generalized version of the above equation includes differential mating choice (non-random mating) and distinction in the fertility of different sexes [Hofbauer and Sigmund, 1998].

### Distortion

Let us now consider the case of distorted allele transmission, a violation of Mendel’s standard segregation law. The gene drives engineered for distortion are in true sense non-Mendelian or super-Mendelian [Goddard and Burt, 1999]. If a drive allele is transmitted from heterozygous parents with probability *p*, the proportion of the three genotypes produced from possible mating pairs can be written as in table 2. The rate of genotype production then changes to

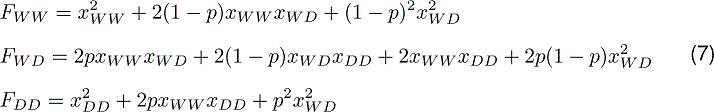

Again the population dynamics for the distorted case is given by Eq. (2), but the effective genotype production rate changes. While deriving Eq. (7) we assume that there is no variation in intrinsic viabilities of the genotypes (*ω* = *ν* = 1), no toxinantidote drive is present (*d_m_* = *d_im_* = *d_s_* = 0) and no fertility selection (*f_WW_* = *f_WD_* = *f_DD_* = 1). We can recover back the standard dynamics for *p* = 0.5 when there is no distortion in transmission probabilities of alleles. If *p >* 0.5, allele transmission from a heterozygote is biased in favour of the driven allele. Heterozygous individuals transmit only the drive allele for *p* = 1. This distortion is also the case of ‘homing drive’ with 100% drive efficiency.

### Combined Dynamics

The rate of the production for the three genotypes because of viability selection, fertility selection and distortion is given by

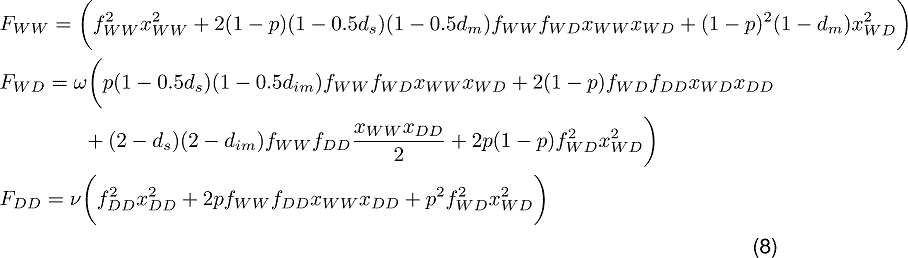

The population dynamics for the combined case is then given by including the above *F_i_*’s in (2).

### Refractory zone

For estimating the refractory zone, we analytically approximated the equation of unstable manifold when distortion and fertility selection both acts at the same time. Setting viability parameters to *ω* = *ν* = 1, no toxin-antidote based drive *d_m_* = *d_im_* = *d_s_* = 0, the rate of offspring production in the next generation is given by:

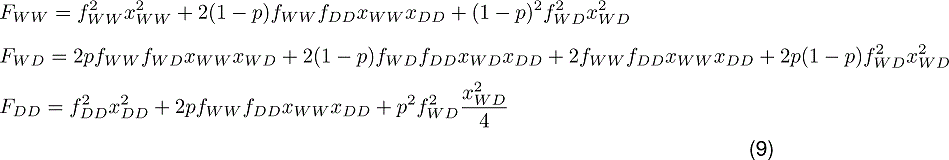

Using the fact that *x_WD_* = 1 *− x_WW_ − x_DD_*, the three population dynamic (2) for the three genotypes can be reduced to two. Keeping all other parameters fixed but *p* and *f_WD_*, we found that an unstable fixed point exists in the interior of the simplex at 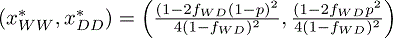. From the chain rule of derivatives, we can write

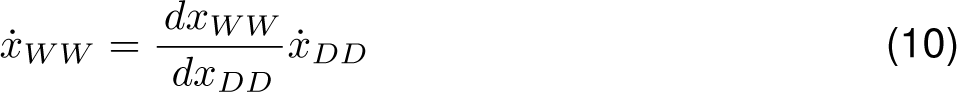

Now, we approximate *x_DD_* by a polynomial of single indeterminate *x_WW_* keeping other parameters constant.

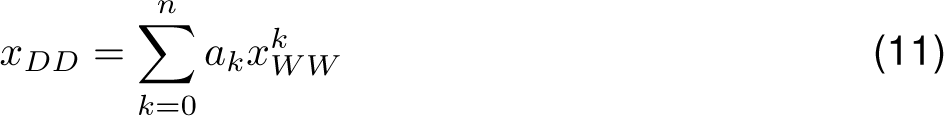

where *a_k_* are the coefficients of the polynomial and *n* has a finite value. Substituting Eq. (11) in Eq. (10) and comparing the coefficients on both sides gives us many solutions for Eq. (11). The correct solution can be filtered by imposing an additional condition that the polynomial passes through the unstable fixed point in the interior of the simplex. Incidentally, the approximated polynomial is a line equation. Finally, the refractory area can be calculated by obtaining the coordinates of the points intersecting the vertex of the simplex. The appropriate codes for the calculations are available on ShinyApps.

### Spatial organization within a population

In this analysis, we use the framework of evolutionary game theory and track the allele frequencies instead of genotype frequencies. The central idea of evolutionary game theory is that the game’s payoff matrix defines the outcome of pairwise interaction between individual entities. Furthermore, the evolutionary success of these individuals is determined by the game’s payoff matrix. In our case, interaction takes place in the allele space, so an individual unit is an allele that can be wildtype (W) of drive type (D). As explored before [Haig, 2010, Traulsen and Reed, 2012] under suitable assumptions, the payoff matrix for meiotic drive, i.e. with distortion and selection is given by:

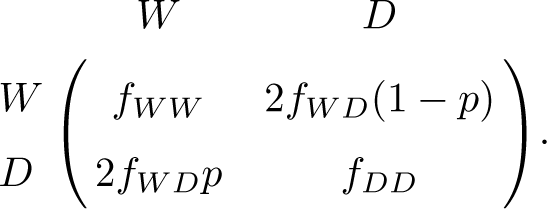

The equation that governs the population dynamics at allele level is then given by the standard selection equation [Crow and Kimura, 1970, Hofbauer and Sigmund, 1998]:

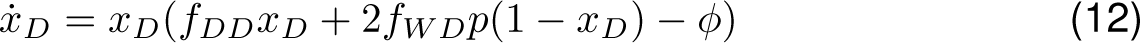

where 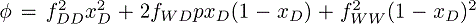 is the average fitness of W and D alleles. The drive allele can invade if *2f_WD_p < f_WW_* (as derived in Noble et al. [2017] and fix in the population if 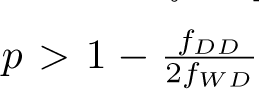. Describing the dynamics using selection equations allows us to write the population dynamics of the gene drive on a regular graph specifically for infinitely large Bethe lattices of degree *k* using the pair-approximation method. Incidentally, this equation is the replicator equation with transformed payoff matrix used in studying evolutionary games on networks [Ohtsuki and Nowak, 2006]. The payoff matrix transformation is different for different update rules. Population update rule defines the elementary process that changes the frequency of each type in the population and usually defined for a finite population. We will use the birth-death update rule in our analysis. In the birth-death update rule, first, an individual is selected proportional to its fitness which then replaces one of its randomly chosen neighbours. Let us consider a game with the payoff matrix *A* = [*a_ij_*] where *i* & *j* can be 1 or 2. Here 1 is wildtype (W) allele and 2 is drive allele (D). When the allele interactions occur on a regular graph of degree *k*, the population dynamics is still represented by the replicator equation but with a transformed payoff matrix. The payoff matrix is transformed to *A,* = [*a_ij_*] + [*b_ij_*] [Ohtsuki and Nowak, 2006] where,

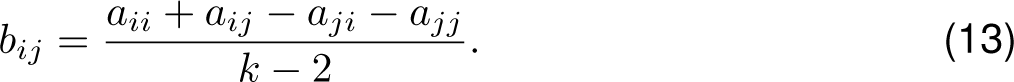

As *k → ∞*, *b_ij_*will become increasingly small, and invasion of gene drive will essentially depend upon whether drive allele is more fit than wildtype when drive is rare, and fixation will depend on whether the same is true when wildtype is rare. The driven gene will invade (from rare) and fix in the population if *a*_21_ + *b*_21_ *> a*_11_ + *b*_11_ and *a*_22_ + *b*_22_ *> a*_12_ + *b*_12_. The conditions for invasion from rarity for the case of distortion and fertility selection is:

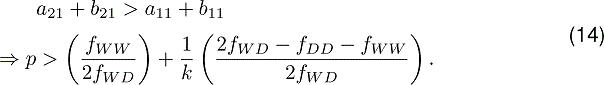

If 2*f_WD_> f_DD_* + *f_WW_*, the critical *p* required for invasion increases relative to the mixed population scenario. Hence a lower network degree *k* results in higher critical *p_c_*. If 2*f_WD_< f_DD_* + *f_WW_*, the critical *p* required for invasion decreases. The condition obtained for the mixed population regime is recovered in the limit of *k → ∞*. The additional condition for the fixation of the gene drive is:

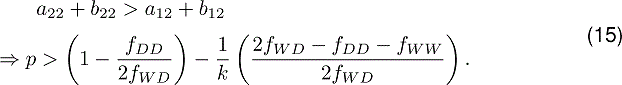

A condition for fixation can be recovered for the mixed population regime in the limit of *k → ∞*. It is also worth noting that the condition for invasion and fixation remains intact with variation in *k* if 2*f_WD_*= *f_DD_* + *f_WW_*. Nevertheless, a constraint exists on the invasion and fixation conditions.

## Acknowledgements

We thank the scientific inputs of Johannes Frieß, Mathias Otto and Samson Simon.

## Availability of data and materials

The appropriate codes for the calculations and literature database are available on GitHub. The deployed app is also available at ShinyApps.

## Authors’ contributions

P.V. and C.S.G. developed the model. All authors conceived the project, developed the theory and wrote the paper.

## Appendix: Backward compatibility

In this section, we will demonstrate the flexibility of our generic modelling approach by recovering the results of earlier work on different gene drive systems. Here we present population dynamics of the three genotypes WW, WD and DD for some special cases using our generic model. Next we show how our base model can be extended to include the possibility of resistance and multi-locus gene drive. Please note that the results shown here are only a subset of the work done in the original studies.

## Recovering Noble et al. Science Advances (2017)

Noble et al. [2017] studied the population dynamics of CRISPR based homing endonuclease gene drive. These gene drive constructs induce a double strand break at the target sequence (wildtype allele). The drive is then copied at the break site using homologous recombination. If resistance evolution is ignored, the final consequence is that the heterozygous individuals only transmit drive allele during recombination. In our generic model, the drive acts in the gamete stage and uses distortion for propagating the drive allele in the population. The authors also accounted for the variation in the fertility rates of genotypes due to the drive construct. Hence every individual undergoes both distortion and fertility selection during its life cycle. We can recover the population dynamics equations for the case using information provided in Table 2 for distortion and fertility selection. The authors derived the following condition which leads to the invasion of wildtype population by the gene drive:

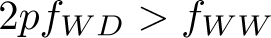

The above invasion condition of Noble et al. [2017] is demonstrated in Fig. A.1. The original study also analyzed the implication of resistance evolution and utility of multiple guide RNAs construct on the evolutionary dynamics. These features can also be included in our model and would entail the addition of more genotypes and their corresponding dynamics.

**Figure A.1:**
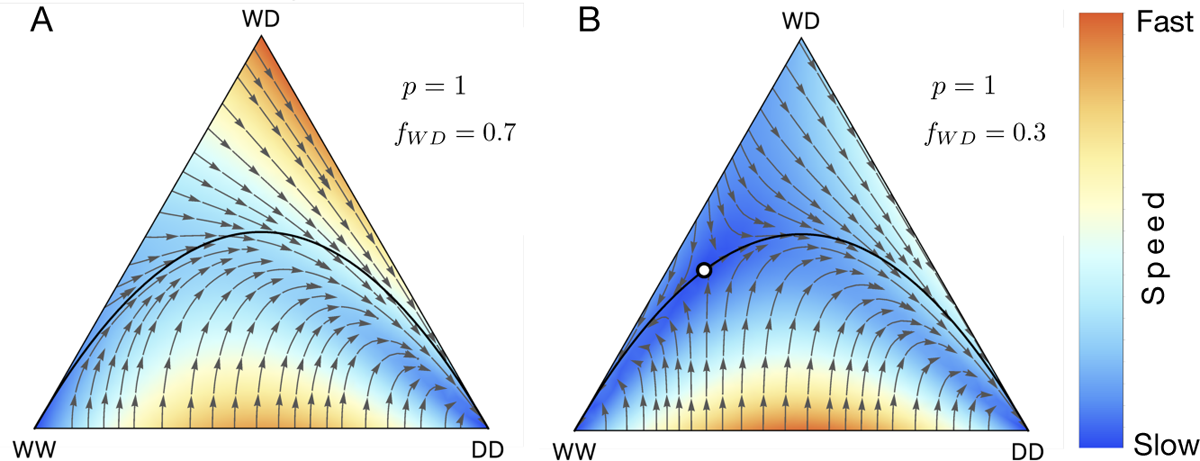
Population dynamics of CRISPR based homing endonuclease gene drive. (A) When the fertility rate of heterozygous adults is 0.7 and drive efficiency is 100%, we have 2*pf_WD_ > f_WW_*. A small release of WD/DD will invade the population consisting entirely of WW. (B) When the fertility rate of heterozygous adults is 0.3, we have 2*pf_WD_ < f_WW_*. Successful invasion by gene drive would require threshold release of WD/DD in the population. The position of the unstable fixed point is (*WW,DD*)= (0.286, 0.354). Other parameters are fixed to *f_WW_* = 1*, f_DD_* = 1 for both A and B.

## Recovering Gokhale et al. BMC Evolutionary Biology (2014)

Gokhale et al. [2014] analysed the synergistic effect of combined Medea and single-locus engineered underdominance in a single transgenic construct. Medea gene drive utilize viability selection which acts during the zygote stage of an organism. In the Medea constructs, wildtype homozygous offspring of a heterozygous mother becomes non-viable (See Table 2C). In single-locus engineered underdominance, the heterozygotes are less fit than both wild and drive homozygotes. Population dynamics of Medea and underdominance can be recovered from Eq. (2) and Eq. (5). Fig. A.2 recovers the results of Gokhale et al. [2014] for special parameter set.

**Figure A.2:**
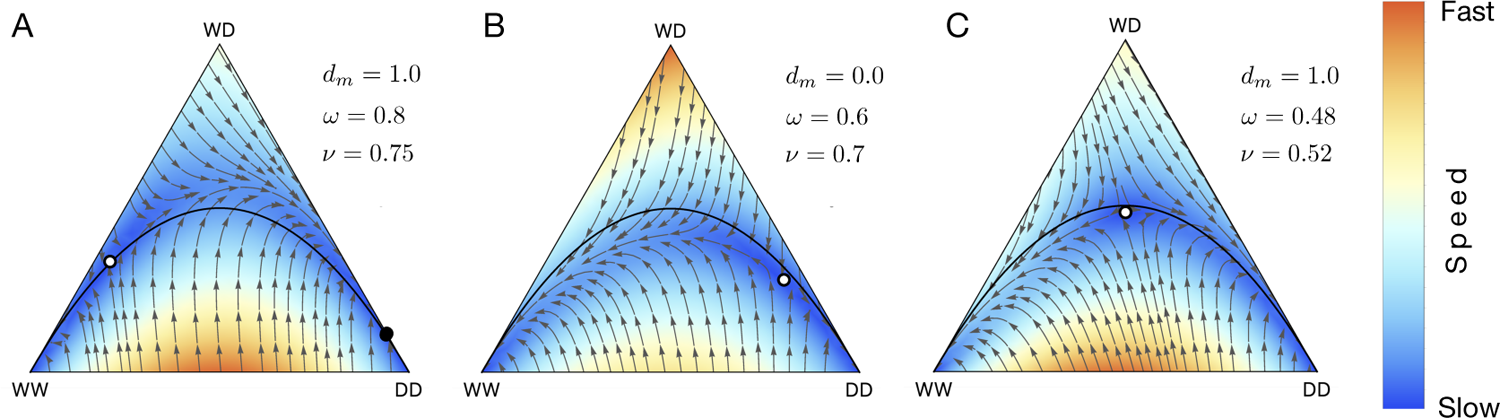
de Finetti diagram showing the population dynamics of Medea, underdominace and their combined effect. (**A**) Medea only (**B**) Underdominance only (**C**) Combined effect of Medea and underdominance

## Recovering Marshall and Hay, Journal of Heredity (2011)

Marshall and Hay [2011] first proposed inverse *Medea* to bring about population replacement but the spread is confined to its released site. In inverse *Medea*, homozygous offspring of a wildtype mother are non-viable (see table A.1).

## Recovering Marshall et al, Genetics (2011)

*Semele* drive was first proposed in Marshall et al. [2011] and is based on toxinantidote system. Transgenic males carry a toxin, and transgenic females carry the corresponding antidote. Offspring of a transgenic male carrying toxin and wildtype female with no antidotes are non-viable. The proportions of offspring of different genotypes is given in table A.1. *Semele* drive like Medea and Inverse Medea utilize viability selection and acts during the zygote stage. The dynamical equation for the minimal case can be recovered using table A.1, are visualised in Fig. A.4.

**Figure A.3:**
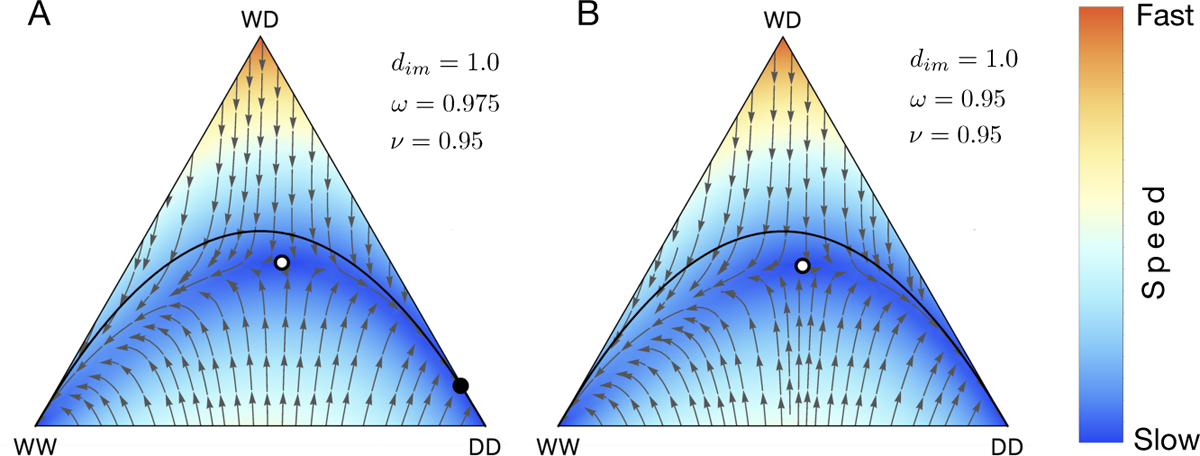
Population dynamics of Inverse Medea. (**A**) For *ω* = 0.975 and *ν* = 0.95 if transgenic individuals are released above a threshold, population converges to a stable point consisting of 99.7% of DD and WD. The stable and unstable fixed point is represented by black and white circle on the de finetti diagram. (**B**) For *ω* = 0.95 and *ν* = 0.95 above a threshold release, drive homozygous (DD) invades the whole population. *dim* = 1

**Figure A.4:**
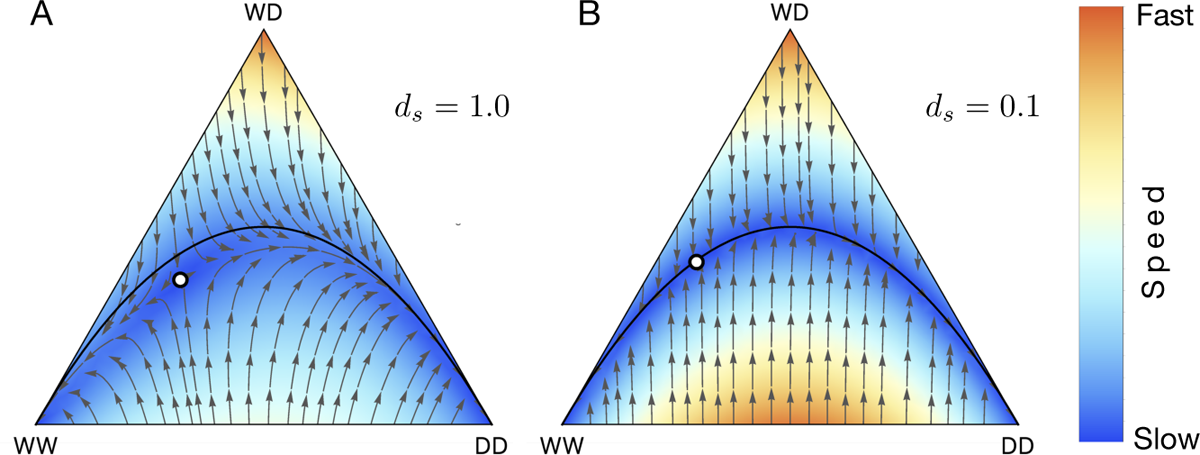
Population dynamics of *Semele* drive when there is no fitness cost. (**A**) Drive efficiency is 100% (**B**) Drive efficiency is 10%

## Resistant allele

Gene drives are prone to resistance evolution due to standing genetic variation or because of the inefficiency of the drive mechanism [Burt, 2003, Esvelt et al., 2014, Deredec et al., 2008]. For example, in CRISPR based homing drives, resistance could arise because the cell repairs the double-stranded break by CRISPR through non-homologous end joining (NHEJ) instead of expected homologous recombination (HR) [Noble et al., 2017]. Many studies have suggested that the drive resistance can severely impact the spread of the gene drive unless mitigating strategies are included [Burt, 2003, Esvelt et al., 2014, Deredec et al., 2008, Noble et al., 2017, Gomulkiewicz et al., 2021, Champer et al., 2018]. Here, we extend our base model to include a drive resistance allele (R). Our mathematical framework is flexible to include the complexity of such resistance evolution in gene drives. It is important to note that these extensions demonstrate our modelling framework’s flexibility to include more complexity. They have not been deployed in the current instance of our `DrMxR` app.

Including an extra allele results in six possible genotype combinations for a single locus diploid population: WW, WD, DD, WR, DR, RR. The table A.2 shows the proportion of different genotypes produced from 36 (6 *×* 6) possible mating pairs. To keep things simpler, we do not show here any fitness variation due to viability or fertility selection and take the example of resistance evolution in CRISPR based homing gene drives. The rate of production of different genotype is given by:

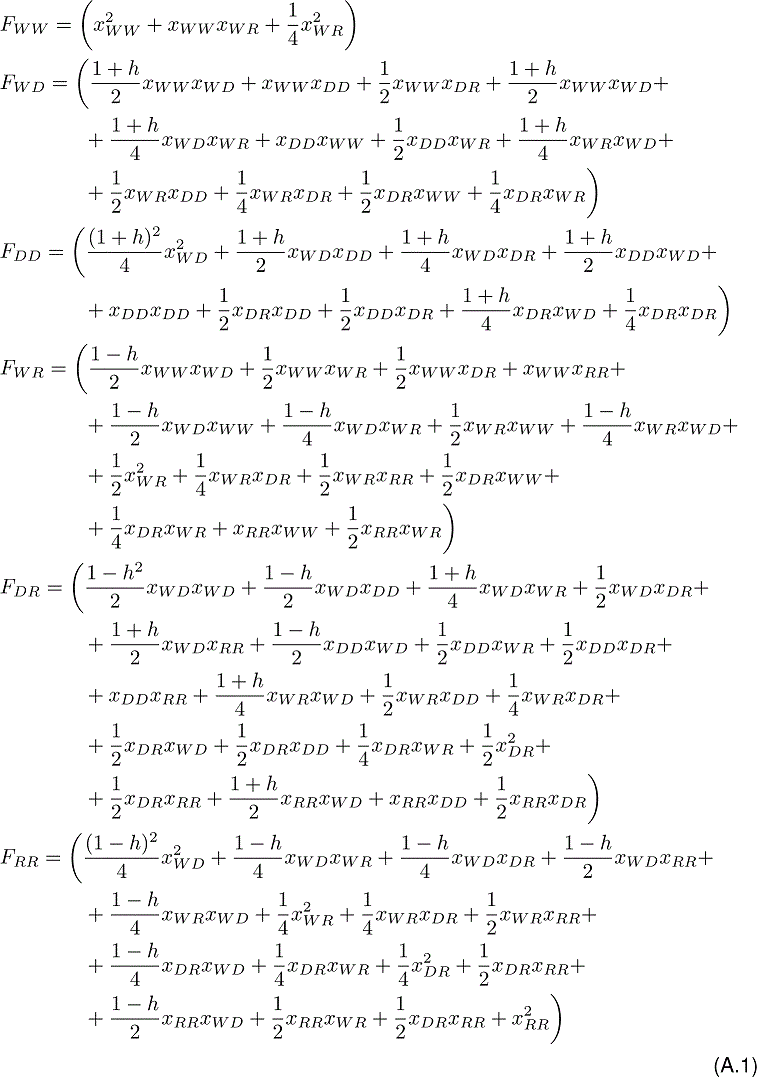

where *h* is the homing efficiency of the CRISPR gene drive hence the probability with which drive heterozygotes parent WD produces gamete with haplotype D and R are 0.5(1 + *h*) and 0.5(1 *−h*) respectively. The population dynamics for the combined case is then given by including the above *F_i_*’s in (2). The resulting dynamical equations are equivalent to the equations obtained by Noble et al. 2017 when there is one resistant allele and no all genotypes have equal fitness [Noble et al., 2017]. The possibility of multiple gRNAs and resistance evolution can also be implemented since the genotype frequencies remain constant:

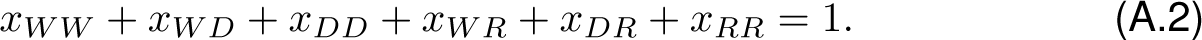

Given the six genotypes, the system’s population dynamics proceeds in a five-dimensional space and cannot be represented in a de Finetti diagram. The specific dynamics could still be studied by numerically solving the equation for various input initial conditions.

## One locus two toxin (1L2T) gene drive

Interestingly, the dynamical equation obtained using Eq.(A.1) demonstrates the addition of multiple alleles to our base model. In this case, the third allele (R) happens to be the resistant allele, but that is not a general case. Like the two allele system, if we remove the distortion because of homing (*h* = 0) and add the effect of fertility or viability selection, the other three allele gene drive systems could be captured through our model. One locus two toxins (1L2T) system is an example of a system where two different drive alleles exist at a single genomic locus like D, and R [Davis et al., 2001, Dhole et al., 2018, Champer et al., 2020c]. The two drive allele, D and R, both encode a different toxin and carry an RNAi (the “antidote”) that neutralizes the other drive allele’s toxin. Therefore, the genotypes containing toxin but no corresponding antidote (WD, RR, DD and WR) are non-viable. In contrast, the viable genotypes are heterozygotes with the two drive alleles (RD) and wild-type homozygotes (WW).

## Multi locus gene drives (Daisy chain drive)

Here we demonstrate that our basic model could be extended to include several multi locus gene drive system [Davis et al., 2001, Dhole et al., 2018, Noble et al., 2019, Champer et al., 2020c]. Daisy chain gene drive is an example of such a drive system [Noble et al., 2019]. It consists of a linear series of genetic elements on different locus where one element drives the next. The last genetic element in the chain is driven to a high frequency, while the element at the base cannot be driven and is lost over time due to natural selection. This process causes the next element to stop driving in the population, and so on. The process continues until the whole population returns to an all wildtype state. Again, owing to plural terminology, the daisy chain system is also referred to as a self-exhausting gene drive [Noble et al., 2019].

To model a multilocus gene drive system, we illustrate a two-locus diploid organism with loci 1 and 2. There are two alleles, the wildtype (W) and the drive type (D). The allele at first loci can therefore be 1*_W_* or 1*_D_*. Similarly, the allele at the second loci is represented by 2*_W_* or 2*_D_*. The genotype corresponding to wildtype homozygous individual at both the loci is 1*_WW_* 2*_WW_*. There are in total nine possible genotypes: 1*WW* 2*WW*, 1*WW* 2*WD*, 1*WW* 2*DD*, 1*WD*2*WW*, 1*WD*2*WD*, 1*WD*2*DD*, 1*DD*2*WW*, 1*DD*2*WD* and 1*_DD_*2*_DD_*. A daisy chain drive uses CRISPR genome editing technology to engineer drive alleles. The drive allele (1*_D_*) in the first locus induces the cutting of the 2*_W_* allele. Considering the nature of distortion outlined in the original paper [Noble et al., 2019], the proportion of offspring from all possible 81 mating pairs can be computed to yield equivalent population dynamic equations [Noble et al., 2019]. A natural extension would be to generalize the framework for any number of locus and allele.

Other multilocus gene drive systems such as two-locus two toxin (2L2T), reciprocal chromosomal translocation (RCT) underdominance system and killer & rescue drive can also be modelled through our framework (if distortion due to homing is not considered). Specific genotype becomes non-viable because of the toxin carrying drive element [Dhole et al., 2018, Champer et al., 2020c]. Besides the wildtype allele, this system consists of two drive alleles at the two loci (say 1*_D_* and 2*_D_*). In reciprocal chromosomal translocation (RCT), the only viable genotypes are homozygotes for the wild-type alleles (1*_WW_*2*_WW_*), homozygotes for the translocated alleles (1*_DD_*2*_DD_*), heterozygotes for the translocated alleles (1*_WD_*2*_WD_*) [Curtis, 1968, Champer et al., 2020c]. While in two locus two toxin (2L2T) system the viable genotypes are homozygotes for the wild-type alleles (1*_WW_*2*_WW_*) and those which carry atleast one copy of each drive allele (1*_WD_*2*_WD_*, 1*_DD_*2*_WD_*, 1*_WD_*2*_DD_*, 1*_DD_*2*_DD_*) [Davis et al., 2001, Champer et al., 2020c]. Killer & rescue gene drive constructs consist of two alleles, namely killer (K) and rescue allele (R), and their corresponding wildtype counterparts are ‘k’, and ‘r’ respectively [Gould et al., 2008]. If the locus of insertion of allele K or R is independent of other loci, there are nine possible genotypes. Out of nine genotype (1*KK* 2*RR*, 1*KK* 2*Rr*, 1*Kk*2*RR*, 1*Kk*2*Rr*, 1*kk*2*RR*, 1*kk*2*Rr*, 1*kk*2*rr*, 1*Kk*2*rr*, and 1*KK* 2*rr*).The genotypes which carry only killer allele K and no rescue allele are non-viable (1*_Kk_*2*_rr_*, and 1*_KK_*2*_rr_*).

Underdominance tethered homing drive (UTH) consist of two components and three alleles with either a transgenic (D) or widtype (W) [Dhole et al., 2019]. This gene drive system can have 27 different diploid genotypes and hence 729 mating possibilities. The details about the fitness of viable and non-viable genotype can be found in the supplementary material of the original study [Dhole et al., 2019]. The wildtype genotype can be represented as 1*_WW_* 2*_WW_* 3*_WW_*. First component is a two-locus engineered underdominance drive which we have already described. The second component is an unlinked locus to be inserted into a haploinsufficient gene, that is, two copies of a functional gene are required at this locus for viable offspring. The homing component at the third locus is driven by the presence of the other two constructs. The guide RNA and Cas endonuclease target the wild-type (3*_W_*) alleles for multiple double-stranded breaks. Repairs through Nonhomologous end-joining (NHEJ) or homology-directed repair (HDR) that did not produce a functional copy of the haploinsufficient results in individuals that are incapable of producing viable offspring. This gene drive system thus helps to prevent the emergence of resistance due to NHEJ [Esvelt et al., 2014].

**Table A.1:** Offspring proportions for Inverse Medea and *Semele* gene drive. Inverse Medea is a two-component system with a zygotic toxin and a maternal antidote. Thus the heterozygous offspring of the wildtype mothers have reduced viability (1 *− dim*). In *Semele* the males carry the toxin while the females carry the antidote. Thus offspring inheriting no antidote from their mothers have reduced viability (1 *− ds*).

| Parents |  | Inverse Medea |  |  | *Semele* |  |  |
| --- | --- | --- | --- | --- | --- | --- | --- |
| ♂ | ♀ | WW | WD | DD | WW | WD | DD |
| WW | WW | 1.0 |  |  | 1.0 |  |  |
| WW | WD | 0.5 | 0.5 |  | 0.5 | 0.5 |  |
| WW | DD |  | 1.0 |  |  | 1.0 |  |
| WD | WW | 0.5 | $0.5(1 - d_{im})$ | | $0.5(1 - d_s)$ | $0.5(1 - d_s)$ | |
| WD | WD | 0.25 | 0.5 | 0.25 | 0.25 | 0.5 | 0.25 |
| WD | DD |  | 0.5 | 0.5 |  | 0.5 | 0.5 |
| DD | WW | | $(1 - d_{im})$ | | | $(1 - d_s)$ | |
| DD | WD |  | 0.5 | 0.5 |  | 0.5 | 0.5 |
| DD | DD |  |  | 1.0 |  |  | 1.0 |

**Table A.2:**
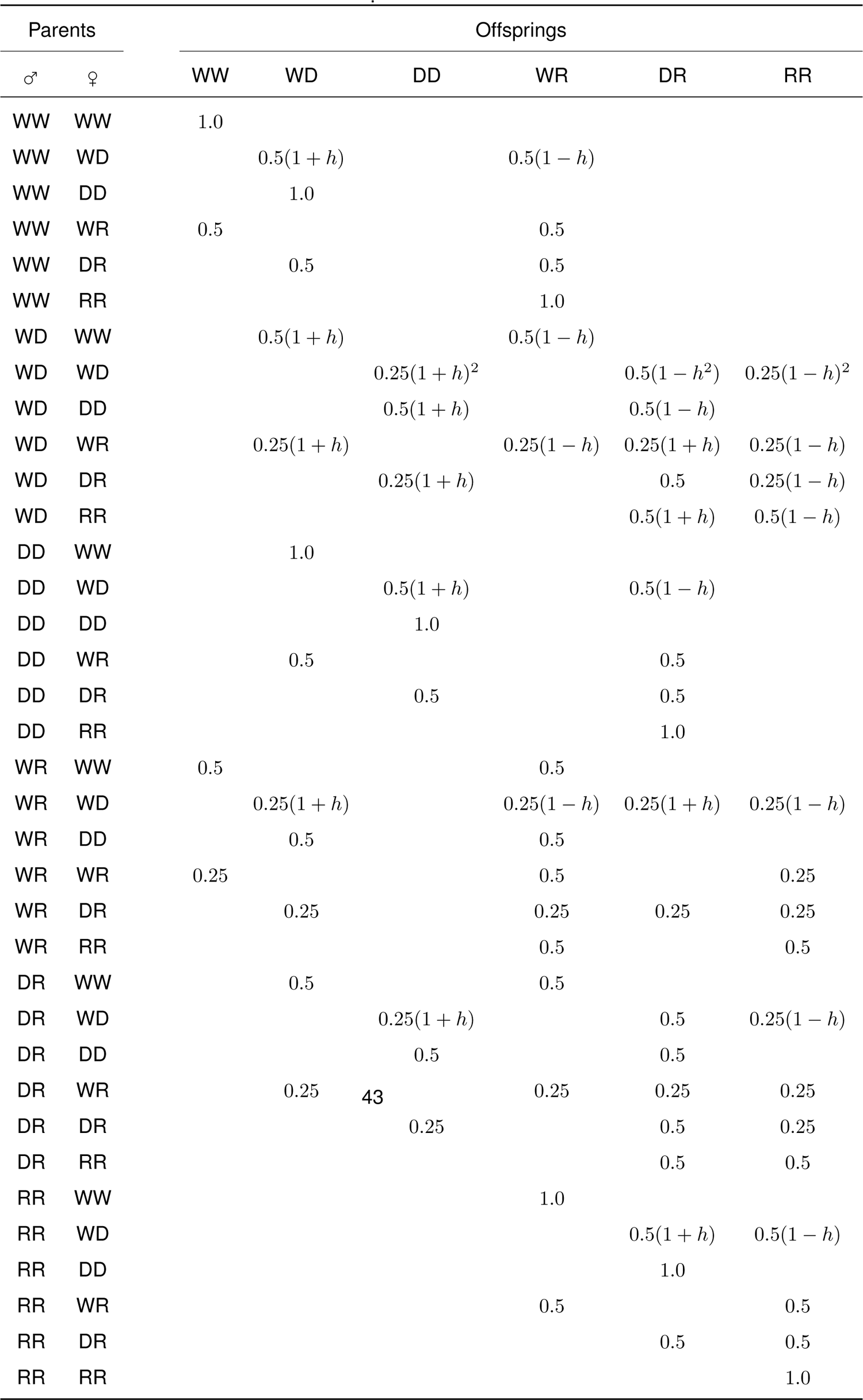
Offspring proportions for CRISPR based homing gene drive with resistance.

## References

1. Omar S Akbari, Kelly D Matzen, John M Marshall, Haixia Huang, Catherine M Ward, and Bruce A Hay. A synthetic gene drive system for local, reversible modification and suppression of insect populations. Current biology, 23(8):671–677, 2013.

2. Omar S Akbari, Chun-Hong Chen, John M Marshall, Haixia Huang, Igor Antoshechkin, and Bruce A Hay. Novel synthetic medea selfish genetic elements drive population replacement in drosophila; a theoretical exploration of medea-dependent population suppression. ACS synthetic biology, 3(12):915–928, 2014.

3. Luke S Alphey, Andrea Crisanti, Filippo Fil Randazzo, and Omar S Akbari. Opinion: Standardizing the definition of gene drive. Proceedings of the National Academy of Sciences, 117(49):30864–30867, 2020.

4. P. M. Altrock, A. Traulsen, R. G. Reeves, and F. A. Reed. Using underdominance to bi-stably transform local populations. Journal of Theoretical Biology, 267:62–75, 2010.

5. P. M. Altrock, A. Traulsen, and F. A. Reed. Stability properties of underdominance in finite subdivided populations. PLoS Computational Biology, 7:e1002260, 2011.

6. Gregory A Backus and Jason A Delborne. Threshold-Dependent Gene Drives in the Wild: Spread, Controllability, and Ecological Uncertainty. BioScience, 69(11):900– 907, 2019.

7. Gregory A Backus and Kevin Gross. Genetic engineering to eradicate invasive mice on islands: modeling the efficiency and ecological impacts. Ecosphere, 7(12):116, 2016.

8. Andrea Beaghton, Pantelis John Beaghton, and Austin Burt. Vector control with driving y chromosomes: modelling the evolution of resistance. Malaria journal, 16(1): 286, 2017.

9. R. W. Beeman, K. S. Friesen, and R. E. Denell. Maternal-effect selfish genes in flour beetles. Science, 256:89–92, 1992.

10. Cara L. Brand, Amanda M. Larracuente, and Daven C. Presgraves. Origin, evolution, and population genetics of the selfish segregation distorter gene duplication in european and african populations of drosophila melanogaster. Evolution, 69(5): 1271–1283, 2015.

11. Anna Buchman, John M Marshall, Dennis Ostrovski, Ting Yang, and Omar S Akbari. Synthetically engineered Medea gene drive system in the worldwide crop pest Drosophila suzukii. Proceedings of the National Academy of Sciences of the United States of America, 115(18):4725–4730, 2018.

12. James J Bull, Christopher H Remien, Richard Gomulkiewicz, and Stephen M Krone. Spatial structure undermines parasite suppression by gene drive cargo. PeerJ, 7: e7921, 2019.

13. Austin Burt. Site-specific selfish genes as tools for the control and genetic engineering of natural populations. Proceedings of the Royal Society B: Biological Sciences, 270(1518):921–928, 2003.

14. Austin Burt and Anne Deredec. Self-limiting population genetic control with sex-linked genome editors. Proceedings. Biological sciences / The Royal Society, 285(1883): 20180776, 2018.

15. Jackson Champer, Anna Buchman, and Omar S Akbari. Cheating evolution: engineering gene drives to manipulate the fate of wild populations. Nature Reviews Genetics, 17(3):146–159, 2016.

16. Jackson Champer, Jingxian Liu, Suh Yeon Oh, Riona Reeves, Anisha Luthra, Nathan Oakes, Andrew G Clark, and Philipp W Messer. Reducing resistance allele formation in crispr gene drive. Proceedings of the National Academy of Sciences, 115 (21):5522–5527, 2018.

17. Jackson Champer, Isabel Kim, Samuel E Champer, Andrew G Clark, and Philipp W Messer. Suppression gene drive in continuous space can result in unstable persistence of both drive and wild-type alleles. bioRxiv, 28:769810, 2019.

18. Jackson Champer, Isabel K Kim, Samuel E Champer, Andrew G Clark, and Philipp W Messer. Performance analysis of novel toxin-antidote crispr gene drive systems. BMC biology, 18(1):1–17, 2020a.

19. Jackson Champer, Esther Lee, Emily Yang, Chen Liu, Andrew G Clark, and Philipp W Messer. A toxin-antidote crispr gene drive system for regional population modification. Nature communications, 11(1):1–10, 2020b.

20. Jackson Champer, Joanna Zhao, Samuel E Champer, Jingxian Liu, and Philipp W Messer. Population dynamics of underdominance gene drive systems in continuous space. ACS Synthetic Biology, 9(4):779–792, 2020c.

21. Jackson Champer, Samuel E. Champer, Isabel K. Kim, Andrew G. Clark, and Philipp W. Messer. Design and analysis of crispr-based underdominance toxinantidote gene drives. Evolutionary Applications, 14(4):1052–1069, 2021.

22. Chun-Hong Chen, Haixia Huang, C. M. Ward, J. T. Su, L. V. Schaeffer, M. Guo, and B. Hay. A synthetic maternal-effect selfish genetic element drives population replacement in drosophila. Science, 316:597–600, 1997.

23. Frank H Collins and Anthony A James. Genetic modification of mosquitoes. Science and Medicine, 3:52–61, 1996.

24. James P Collins. Gene drives in our future: challenges of and opportunities for using a self-sustaining technology in pest and vector management. BMC Proceedings, 12(S8):9, 2018. ISSN 1753-6561.

25. Virginie Courtier-Orgogozo, Baptiste Morizot, and Christophe Boete. Agricultural pest control with crispr-based gene drive: time for public debate: should we use gene drive for pest control? EMBO reports, 18(6):878–880, 2017.

26. GB Craig, WA Hickey, and RC VandeHey. An inherited male-producing factor in aedes aegypti. Science, 132(3443):1887–1889, 1960.

27. J. F. Crow. Why is mendelian segregation so exact? BioEssays, 13:305–312, 1991.

28. J. F. Crow and M. Kimura. *An Introduction to Population Genetics Theory*. Harper and Row, New York, 1970.

29. C.F. Curtis. Possible use of translocations to fix desirable genes in insect pest populations. Nature, 218:368–369, 1968.

30. S. Davis, N. Bax, and P. Grewe. Engineered underdominance allows efficient and economical introgression of traits into pest populations. Journal of Theoretical Biology, 7:83–98, 2001.

31. Anne Deredec, Austin Burt, and H Charles, J Godfray. The population genetics of using homing endonuclease genes in vector and pest management. Genetics, 179 (4):2013–2026, 2008.

32. Sumit Dhole, Michael R Vella, Alun L Lloyd, and Fred Gould. Invasion and migration of spatially self-limiting gene drives: A comparative analysis. Evolutionary Applications, 11(5):794–808, 2018.

33. Sumit Dhole, Alun L Lloyd, and Fred Gould. Tethered homing gene drives: a new design for spatially restricted population replacement and suppression. Evolutionary applications, 12(8):1688–1702, 2019.

34. Sumit Dhole, Alun L Lloyd, and Fred Gould. Gene drive dynamics in natural populations: The importance of density-dependence, space and sex. arXiv, 2020.

35. James E DiCarlo, Alejandro Chavez, Sven L Dietz, Kevin M Esvelt, and George M Church. Safeguarding crispr-cas9 gene drives in yeast. Nature biotechnology, 33 (12):1250, 2015.

36. Kelly A Dyer and David W Hall. Fitness consequences of a non-recombining sex-ratio drive chromosome can explain its prevalence in the wild. Proceedings of the Royal Society B, 286(1917):20192529, 2019.

37. Philip A Eckhoff, Edward A Wenger, H Charles, J Godfray, and Austin Burt. Impact of mosquito gene drive on malaria elimination in a computational model with explicit spatial and temporal dynamics. Proceedings of the National Academy of Sciences, 114(2):E255–E264, 2017.

38. Matthew P Edgington and Luke S Alphey. Population dynamics of engineered underdominance and killer-rescue gene drives in the control of disease vectors. PLoS computational biology, 14(3):e1006059, 2018.

39. Matthew P Edgington and Luke S Alphey. Modeling the mutation and reversal of engineered underdominance gene drives. Journal of theoretical biology, 479:14– 21, 2019.

40. Matthew P Edgington, Tim Harvey-Samuel, and Luke Alphey. Split drive killer-rescue provides a novel threshold-dependent gene drive. Scientific reports, 10(1):1–13, 2020.

41. Kevin M Esvelt, Andrea L Smidler, Flaminia Catteruccia, and George M Church. Concerning RNA-guided gene drives for the alteration of wild populations. eLife, 3: 20131071, 2014.

42. Nicky R Faber, Gus R McFarlane, R Chris Gaynor, Ivan Pocrnic, C Bruce A, Whitela w, and Gregor Gorjanc. Novel combination of crispr-based gene drives eliminates resistance and localises spread. Scientific reports, 11(1):1–15, 2021.

43. Marcus W Feldman and Uri Liberman. A symmetric two-locus fertility model. Genetics, 109(1):229–253, 1985.

44. Sam Ronan Finnegan, Nathan Joseph White, Dixon Koh, M Florencia Camus, Kevin Fowler, and Andrew Pomiankowski. Meiotic drive reduces egg-to-adult viability in stalk-eyed flies. Proceedings of the Royal Society B, 286(1910):20191414, 2019.

45. Johannes L Frieß, Arnim von Gleich, and Bernd Giese. Gene drives as a new quality in gmo releases—a comparative technology characterization. PeerJ, 7:e6793, 2019.

46. V M Gantz, N Jasinskiene, O Tatarenkova, A Fazekas, V M Macias, E Bier, and A A James. Highly efficient cas9-mediated gene drive for population modification of the malaria vector mosquito anopheles stephensi. Proceedings of the National Academy of Sciences, 112(49):E6736–E6743, 2015.

47. Valentino M Gantz and Ethan Bier. The mutagenic chain reaction: a method for converting heterozygous to homozygous mutations. Science, 348(6233):442–444, 2015.

48. Leo Girardin, Vincent Calvez, and Florence Debarre. Catch Me If You Can: A Spatial Model for a Brake-Driven Gene Drive Reversal. Bulletin of Mathematical Biology, 81(12):5054–5088, 2019.

49. Matthew R Goddard and Austin Burt. Recurrent invasion and extinction of a selfish gene. Proceedings of the National Academy of Sciences, 96(24):13880–13885, 1999.

50. H Charles, J Godfray, Ace North, and Austin Burt. How driving endonuclease genes can be used to combat pests and disease vectors. BMC biology, 15(1):1–12, 2017.

51. Chaitanya S Gokhale, Richard G Reeves, and F A Reed. Dynamics of a combined medea-underdominant population transformation system. BMC Evolutionary Biology, 14(1):98, 2014.

52. Richard Gomulkiewicz, Micki L Thies, and James J Bull. Evading resistance to gene drives. Genetics, 217(2), 2021.

53. Fred Gould, Yunxin Huang, Mathieu Legros, and Alun L Lloyd. A killer–rescue system for self-limiting gene drive of anti-pathogen constructs. Proceedings of the Royal Society B: Biological Sciences, 275(1653):2823–2829, 2008.

54. Hannah A Grunwald, Valentino M Gantz, Gunnar Poplawski, Xiang-Ru S Xu, Ethan Bier, and Kimberly L Cooper. Super-mendelian inheritance mediated by crispr– cas9 in the female mouse germline. Nature, 566(7742):105, 2019.

55. David Haig. Games in Tetrads: Segregation, Recombination, and Meiotic Drive. The American Naturalist, 176(4):404–413, 2010.

56. Benjamin C Haller and Philipp W Messer. SLiM 3: Forward Genetic Simulations Beyond the Wright-Fisher Model. Molecular biology and evolution, 36(3):632–637, 2019.

57. A Hammond, R Galizi, K Kyrou, A Simoni, C Siniscalchi, D Katsanos, M Gribble, D Baker, E Marois, S Russell, A Burt, N Windbichler, A Crisanti, and T Nolan. A crispr-cas9 gene drive system targeting female reproduction in the malaria mosquito vector anopheles gambiae. Nature Biotechnology, 34:78–83, 2016.

58. D L Hartl. Genetic dissection of segregation distortion ii. mechanism of suppression of distortion by certain inversions. Genetics, 80(3):539–547, 1975.

59. Yuichiro Hiraizumi and Anita M Thomas. Suppressor systems of segregation distorter (sd) chromosomes in natural populations of drosophila melanogaster. Genetics, 106(2):279–292, 1984.

60. J. Hofbauer and K. Sigmund. Evolutionary Games and Population Dynamics. Cam-bridge University Press, Cambridge, UK, 1998.

61. J. Hofbauer, P. Schuster, and K. Sigmund. Game dynamics in mendelian populations. Biological Cybernetics, 43:51–57, 1982.

62. Luke Holman. Evolutionary simulations of z-linked suppression gene drives. Proceedings of the Royal Society B, 286(1912):20191070, 2019.

63. Yunxin Huang, Alun L Lloyd, Mathieu Legros, and Fred Gould. Gene-drive into insect populations with age and spatial structure: A theoretical assessment. Evolutionary applications, 4(3):415–428, 2011.

64. Alison T Isaacs, Fengwu Li, Nijole Jasinskiene, Xiaoguang Chen, Xavier Nirmala, Osvaldo Marinotti, Joseph M Vinetz, and Anthony A James. Engineered resistance to plasmodium falciparum development in transgenic anopheles stephensi. PLoS Pathog, 7(4):e1002017, 2011.

65. Kyros Kyrou, Andrew M Hammond, Roberto Galizi, Nace Kranjc, Austin Burt, Andrea K Beaghton, Tony Nolan, and Andrea Crisanti. A crispr–cas9 gene drive targeting doublesex causes complete population suppression in caged anopheles gambiae mosquitoes. Nature biotechnology, 36(11):1062, 2018.

66. William Larner, Tom Price, Luke Holman, and Nina Wedell. An x-linked meiotic drive allele has strong, recessive fitness costs in female drosophila pseudoobscura. Proceedings of the Royal Society B, 286(1916):20192038, 2019.

67. Amanda M Larracuente and Daven C Presgraves. The selfish segregation distorter gene complex of drosophila melanogaster. Genetics, 192(1):33–53, 2012.

68. Anna K Lindholm, Kerstin Musolf, Andrea Weidt, and Barbara Konig. Mate choice for genetic compatibility in the house mouse. Ecology and evolution, 3(5):1231–1247, 2013.

69. Anna K Lindholm, Kelly A Dyer, Renee C Firman, Lila Fishman, Wolfgang Forstmeier, Luke Holman, Hanna Johannesson, Ulrich Knief, Hanna Kokko, Amanda M Larracuente, et al. The ecology and evolutionary dynamics of meiotic drive. Trends in ecology & evolution, 31(4):315–326, 2016.

70. Mary F Lyon. Transmission ratio distortion in mice. Annual review of genetics, 37(1): 393–408, 2003.

71. John M Marshall. The effect of gene drive on containment of transgenic mosquitoes. Journal of theoretical biology, 258(2):250–265, 2009.

72. John M Marshall and Omar S Akbari. Gene drive strategies for population replacement. In Genetic control of malaria and dengue, chapter 9, pages 169–200. Elsevier, 2016.

73. John M Marshall and Bruce A Hay. Inverse Medea as a Novel Gene Drive System for Local Population Replacement A Theoretical Analysis. Journal of Heredity, 103(3): 336–341, 2011.

74. John M Marshall and Bruce A Hay. Confinement of gene drive systems to local populations: a comparative analysis. Journal of Theoretical Biology, 294:153–171, 2012.

75. John M Marshall, Geoffrey W Pittman, Anna B Buchman, and Bruce A Hay. Semele: a killer-male, rescue-female system for suppression and replacement of insect disease vector populations. Genetics, 187(2):535–551, 2011.

76. John Min, Charleston Noble, Devora Najjar, and Kevin M Esvelt. Daisy quorum drives for the genetic restoration of wild populations. BioRxiv, page 115618, 2017.

77. Dorian Moro, Margaret Byrne, Malcolm Kennedy, Susan Campbell, and Mark Tizard. Identifying knowledge gaps for gene drive research to control invasive animal species: the next crispr step. Global Ecology and Conservation, 13:e00363, 2018.

78. Thomas Nagylaki. Evolution under fertility and viability selection. Genetics, 115(2): 367–375, 1987.

79. Charleston Noble, Jason Olejarz, Kevin M Esvelt, George M Church, and Martin A Nowak. Evolutionary dynamics of CRISPR gene drives. Science Advances, 3(4), 2017.

80. Charleston Noble, Ben Adlam, George M Church, Kevin M Esvelt, and Martin A Nowak. Current crispr gene drive systems are likely to be highly invasive in wild populations. Elife, 7:e33423, 2018.

81. Charleston Noble, John Min, Jason Olejarz, Joanna Buchthal, Alejandro Chavez, Andrea L Smidler, Erika A DeBenedictis, George M Church, Martin A Nowak, and Kevin M Esvelt. Daisy-chain gene drives for the alteration of local populations. Proceedings of the National Academy of Sciences, 116(17):8275–8282, 2019.

82. Ace North, Austin Burt, and H Charles, J Godfray. Modelling the spatial spread of a homing endonuclease gene in a mosquito population. Journal of Applied Ecology, 50(5):1216–1225, 2013.

83. Ace R North and H Charles, J Godfray. The dynamics of disease in a metapopulation: The role of dispersal range. Journal of theoretical biology, 418:57–65, 2017.

84. Ace R North, Austin Burt, and H Charles, J Godfray. Modelling the potential of genetic control of malaria mosquitoes at national scale. BMC biology, 17(1):1–12, 2019.

85. Ace R North, Austin Burt, and H Charles, J Godfray. Modelling the suppression of a malaria vector using a crispr-cas9 gene drive to reduce female fertility. BMC biology, 18(1):1–14, 2020.

86. Georg Oberhofer, Tobin Ivy, and Bruce A Hay. Cleave and rescue, a novel selfish genetic element and general strategy for gene drive. Proceedings of the National Academy of Sciences, 116(13):6250–6259, 2019.

87. Georg Oberhofer, Tobin Ivy, and Bruce A Hay. Gene drive and resilience through renewal with next generation cleave and rescue selfish genetic elements. Proceedings of the National Academy of Sciences, 117(16):9013–9021, 2020.

88. H. Ohtsuki and M. A. Nowak. The replicator equation on graphs. Journal of Theoretical Biology, 243:86–97, 2006.

89. M. F. Palopoli and C. I. Wu. Rapid evolution of a coadapted gene complex: evidence from the segregation distorter (sd) system of meiotic drive in drosophila melanogaster. Genetics, 143:1675–1688, 1996.

90. Tom AR Price and Nina Wedell. Selfish genetic elements and sexual selection: their impact on male fertility. Genetica, 132(3):295, 2008.

91. Thomas AA Prowse, Fatwa Adikusuma, Phillip Cassey, Paul Thomas, and Joshua V Ross. A y-chromosome shredding gene drive for controlling pest vertebrate populations. Elife, 8:e41873, 2019.

92. R. G. Reeves, J. Bryk, P. M. Altrock, J. A. Denton, and F A Reed. First steps towards underdominant genetic transformation of insect populations. PLoS ONE, 9 (5), 2014.

93. Hector M, Sanchez C, Sean L Wu, Jared B Bennett, and John M Marshall. MGDrivE: A modular simulation framework for the spread of gene drives through spatially explicit mosquito populations. Methods in Ecology and Evolution, 11(2):229–239, 2019.

94. L. Sandler and Kent Golic. Segregation distortion in drosophila. Trends in Genetics, 1(C):181–185, 1985.

95. L. Sandler and E. Novitski. Meiotic drive as an evolutionary force. The American Naturalist, 91:105–110, 1957.

96. L Sandler, Yuichiro Hiraizumi, and Iris Sandler. Meiotic drive in natural populations of drosophila melanogaster. i. the cytogenetic basis of segregation-distortion. Genetics, 44(2):233, 1959.

97. Alekos Simoni, Andrew M Hammond, Andrea K Beaghton, Roberto Galizi, Chrysanthi Taxiarchi, Kyros Kyrou, Dario Meacci, Matthew Gribble, Giulia Morselli, Austin Burt, et al. A male-biased sex-distorter gene drive for the human malaria vector anopheles gambiae. Nature biotechnology, 38:1054–1060, 2020.

98. Hidenori Tanaka, Howard A Stone, and David R Nelson. Spatial gene drives and pushed genetic waves. Proceedings of the National Academy of Sciences, 114 (32):8452–8457, 2017.

99. A. Traulsen and F. A. Reed. From genes to games: Cooperation and cyclic dominance in meiotic drive. Journal of Theoretical Biology, 299:120–125, 2012.

100. A. Traulsen, J. C. Claussen, and C. Hauert. Coevolutionary dynamics: From finite to infinite populations. Physical Review Letters, 95:238701, 2005.

101. A. Traulsen, J. C. Claussen, and C. Hauert. Coevolutionary dynamics in large, but finite populations. Physical Review E, 74:011901, 2006.

102. Robert L Unckless, Andrew G Clark, and Philipp W Messer. Evolution of Resistance Against CRISPR/Cas9 Gene Drive. Genetics, 205(2):827–841, 2017.

103. M. van Veelen. Hamilton’s missing link. Journal of Theoretical Biology, 246:551–554, 2007.

104. Michael R Vella, Christian E Gunning, Alun L Lloyd, and Fred Gould. Evaluating strategies for reversing crispr-cas9 gene drives. Scientific reports, 7(1):11038, 2017.

105. M. J. Wade and R. W. Beeman. The population dynamics of maternal-effect selfish genes. Genetics, 138:1309–1314, 1994.

106. Catherine M Ward, Jessica T Su, Yunxin Huang, Alun L Lloyd, Fred Gould, and Bruce A Hay. Medea selfish genetic elements as tools for altering traits of wild populations: a theoretical analysis. Evolution, 65(4):1149–1162, 2011.

107. Kelsey Lane Warmbrod, Amanda Kobokovich, Rachel West, Georgia Ray, Marc Trotochaud, and Michael Montague. Gene drives: Pursuing opportunities, minimizing risk. Report, Johns Hopkins Center for Health Security, May 2020.

108. Katie Willis and Austin Burt. Double drives and private alleles for localised population genetic control. PLoS Genetics, 17(3):e1009333, 2021.

109. Nikolai Windbichler, Miriam Menichelli, Philippos Aris Papathanos, Summer B Thyme, Hui Li, Umut Y Ulge, Blake T Hovde, David Baker, Raymond J Monnat, Austin Burt, and Andrea Crisanti. A synthetic homing endonuclease-based gene drive system in the human malaria mosquito. Nature, 473(7):212–215, 2011.

